# GARD: Genomic Data based Drug Repurposing in Head and Neck Cancer with Large Language Model Validation

**DOI:** 10.64898/2026.01.15.699561

**Authors:** Pradham Tanikella, Will Nenad, Christophe Courtine, Yifan Dai, Qingying Deng, Baiming Zou, Nosayaba Osazuwa-Peters, Travis Schrank, Di Wu

**Affiliations:** Department of Genetics, University of North Carolina at Chapel Hill School of Medicine, Chapel Hill, North Carolina; Lineberger Comprehensive Cancer Center, University of North Carolina at Chapel Hill School of Medicine, Chapel Hill, North Carolina; Department of Otolaryngology/Head and Neck Surgery, University of North Carolina at Chapel Hill School of Medicine, Chapel Hill, North Carolina; Department of Biostatistics, Gillings School of Global Public Health at University of North Carolina at Chapel Hill; Department of Biomedical Sciences, Adams School of Dentistry, University of North Carolina at Chapel Hill; Department of Head and Neck Surgery & Communication Sciences, Duke University School of Medicine; Department of Population Health Sciences, Duke University School of Medicine

**Keywords:** Head and Neck Cancer, Drug repurposing, Genomics

## Abstract

**Background/Objectives:** Head and neck cancer (HNC) represents the seventh most common cancer diagnosis globally, yet current treatments, including surgery, radiation, and immunotherapy, have shown limited improvement in outcomes. Drug repurposing offers a cost-effective strategy to identify new therapeutic options by leveraging existing medications with known safety profiles. Within this study we developed the **GARD** pipeline (**G**enomic **A**lteration-based **R**epurposing for **D**rugs), designed to uncover repurposing candidates for HNC using genomic and network-based approaches.

**Methods:** GARD integrates multi-omics data from The Cancer Genome Atlas (TCGA), including copy number variation (CNV) and somatic mutations (SOM). The cohort was stratified by human papillomavirus (HPV) status. Risk-associated genes were identified and then expanded via high-confidence protein–protein interaction (PPI) networks. Top candidate genes were filtered through comprehensive analysis of publicly available literature data in PubMed using LLM to validate the relationship between the identified genes and HNC. The validated top risk genes and their network-expanded neighbors were mapped against DrugBank, and through statistical significance testing, established significant drug-gene associations.

**Results:** Significant genes associated with HNC, inferred by genomics alteration, were identified across HPV-positive and HPV-negative subgroups, such as PIK3CA, SOX2, TP53, EIF4G1, TLR7, CLDN1, PRKCI, and EPHA2. Further expansion through the PPI network identified other targetable genes such as EGFR, ERBB2, and the FGFRs. Literature based validation efforts ensured provided confidence in the gene-disease association. Drug-gene mapping revealed candidates spanning already in clinical trials for HNC (e.g. Afatinib, Cabozantinib, Dasatinib, Brigatinib, Lenvatinib, Capivasertib, Erdafitinib) and emerging or repurposing candidates (Amuvatinib, XL765 (Voxtalisib), Golotimod, Artenimol, Quercetin, and Acetylsalicylic Acid), offering opportunities for precision repurposing.

**Conclusions:** The GARD pipeline demonstrates a genomics-driven, network-informed framework for systematic drug repurposing in HNC. HPV stratification enhances precision, literature-based validation strengthens confidence, and integrated drug mapping enables refinement of existing therapies and discovery of novel candidates for personalized treatment strategies.

**Code Availability:** The full implementation of the GARD pipeline, including preprocessing scripts, statistical analysis modules, and visualization tools, is publicly available on GitHub at: https://github.com/pvtanike/Genomic-Landscape-Based-Drug-Repurposing.git

**Simple Summary:** Head and neck cancer (HNC) is among the most prevalent and challenging cancers world-wide. Developing new drugs is expensive and time-consuming, so this study explored a faster, cost-effective approach utilizing existing medications with established safety profiles: drug repurposing. We developed the **GARD** pipeline (**G**enomic **A**lteration-based **R**epurposing for **D**rugs), which utilizes large-scale genomic data from The Cancer Genome Atlas (TCGA) to identify key genomics changes in HNC. These genes are expanded through protein–protein interaction networks to capture related pathways and then validated using evidence from thousands of PubMed articles extracted by large language model (LLM) tools. Finally, validated genes are matched with drugs using the DrugBank database. This approach uncovered both known cancer drugs and promising new candidates. These included targeted therapies such as Fostamatinib, Nintedanib, Brigatinib, Regorafenib, and Lenvatinib, as well as emerging compounds like Artenimol, Quercetin, and Acetylsalicylic Acid (Aspirin). Through a combination of genomic analysis, network expansion, and literature validation, the GARD pipeline offers a powerful way to accelerate personalized cancer treatments while reducing cost and development time.

## 1. Introduction

Head and neck cancer (HNC) includes a diverse group of malignancies arising in the oral cavity, pharynx, hypopharynx, larynx, nasal cavity, and other regions of the head and neck, ranking as the seventh most common cancer globally [1]. Common treatments include surgical resection followed by radiotherapy and chemotherapy [2]. The medications/drugs in use today to treat HNC are still limited. Targeted therapies such as kinase inhibitors, and immunotherapies, have shown promise in other cancers and have been used in HNC recently [3]. Despite these advances, the treatment of HNC is still challenging due to the heterogeneity and novel treatments still have a hard time making it to market [2]. Additionally, a crucial factor to consider in HNC is HPV status, which significantly influences tumor biology, treatment response, and overall prognosis. Stratifying patients by HPV status enables more precise identification of genomic alterations and improves the relevance of therapeutic candidates for HPV-positive and HPV-negative subtypes [2,4].

Identifying more targeted treatments could significantly improve outcomes for patients with HNC. One promising and cost-effective strategy is drug repurposing, which involves finding new therapeutic uses for existing medications [5]. This strategy offers a cost-effective and time-efficient alternative to traditional drug development, as repurposed drugs have already undergone extensive safety and pharmacokinetic testing [6]. Computational repurposing approaches, which leverage large-scale biomedical datasets to systematically uncover mechanism-based drug-disease relationships, have shown potential [5–8]. Genomic based computational methods include integration of genomic, transcriptomic, proteomic, and pharmacological data to identify drugs with potential efficacy against molecular targets. By harnessing publicly available multi-omics data, researchers can expand therapeutic options through the repositioning of approved or investigational compounds, ultimately advancing precision oncology [9,10].

In cancer research, genomic based computational drug repurposing has already led to promising therapeutic advances. Recent advancements in available large scale cancer genomics data have allowed for investigation of cancer driving genetic therapeutic targets [11–13]. Cancer is driven by changes in the genome, studying these changes in HNC can help identify key driver genes in tumor initiation and progression [2,14]. In this way, repurposed drug candidates that target key disease-related genes can be identified. The Cancer Genome Atlas (TCGA) and Cancer Cell Line Encyclopedia (CCLE) are some of the datasets that serve as sources for cancer-related genes. They provide information about copy number variations (CNVs), somatic mutations (SOMs), clinical data, and drug screens on cell lines [12,15,16].

TCGA dataset serves as one of the main data source for cancer-related genes. Existing TCGA based drug repurposing varied in approaches and targets. Many studies focused on identifying therapeutic candidates in varied cancers, through differential gene expression, survival-associated markers, or Cox regression models to link genomic features with patient outcomes [17–19]. Some focused on general underlying mechanisms and potential for use in treatment [20,21]. Within HNC specifically, some studies narrowed in on specific medications or niche conditions [22]. Others maintained a broad approach to HNC alongside network expansion as in this study but did not incorporate HPV stratification and chose other methods of validation to land on very few specific repurposing candidates [23]. Some included HPV stratification, but were focused specifically on known or immediate alterations within the HNC genomic landscape and did not incorporate any network expansion [24].

Based on previous repurposing studies, a major limitation of current approaches lies in the lack of rigorous filtering and validation to prioritize a high-confidence set of drugs for clinical application [5]. Current works were able to identify possible repurposing candidates, but utilized one data source and needed larger manual review for identifying top candidates [5,15]. Additionally, many previous studies were effective in drug repurposing but had limited stratification of HPV analysis, or focused on survival-associated genes rather than a comprehensive genomic analysis, and only a few incorporated the use of robust validation steps to confirm candidate relevance and narrow repurposing candidates. The incorporation of network expansion to broadly identify possible HNC targetable genes alongside HPV stratified TCGA data with external validation provides a unique opportunity and approach to producing repurposing candidates.

Uniquely, this study introduces the **GARD** pipeline (**G**enomic **A**lteration-based **R**epurposing for **D**rugs) which integrates HPV stratification with large scale multi-omics to identify key disease related genes within HPV +/- patients. These genes are expanded using high confidence interactions with other proteins and rigorously validated using thousands of peer reviewed articles from PubMed. Lastly identification of medications is completed using DrugBank where known drug-gene combinations are curated and thorough empirical validation is performed. The results from our drug repurposing pipeline, GARD, confirmed the validity of its performance through identification of both established and novel therapeutic candidates for HNC. In this study, we demonstrate that genomics-based computational drug repurposing can provide a scalable and precise framework for discovering new treatment options tailored to molecular subtypes, such as HPV-positive and HPV-negative HNC. Given the heterogeneity of many cancer types and the growing availability of genomic datasets, our pipeline, GARD, holds promise for broader application across other malignancies and even non-cancer diseases, potentially accelerating the discovery of novel therapeutics, reduce development costs, and expand precision medicine.

## 2. Materials and Methods

### 2.1. GARD Pipeline Overview

To identify drug repurposing candidates for HNC, we developed the GARD pipeline. This framework integrates multiple publicly available biomedical resources to combine genomic insights with pharmacological and publicly available literature data for the identification of drug repurposing candidates in HNC.

First, at its core, GARD leverages TCGA, which provides multi-omics datasets, including Copy Number Variation (CNV) and Somatic Mutation (SOM) profiles with associated clinical annotations. The datasets were analyzed to produce high-confidence risk genes associated with HNC. All data was used alongside results of previous work to stratify patients by human papillomavirus (HPV) status [13]. This stratification enables subtype-specific analysis and improves therapeutic relevance [12].

Second, to capture pathway-level context, GARD expands high-confidence risk genes through STRING protein–protein interaction (PPI) networks, incorporating immediate neighbors with strong biological connectivity [25]. These genes are validated through external data sources of publicly available literature data within PubMed, as literature-based repurposing efforts have shown effectiveness [26,27].

Third, after extraction and cleaning, the literature data was parsed by the LLM model, Google GEMMA, to identify genes associated with HNC and provide peer-reviewed evidence as support for the gene association with HNC and filter the identified gene list before input into DrugBank. [28].

Fourth, to connect these validated genetic findings with potential therapeutic agents, GARD incorporates curated drug–gene data from DrugBank [29]. To ensure the that drugs identified have a significant relationship with target genes identified either directly from CNV/mutation analysis or as the protein neighbor genes, statistical enrichment-based tests were performed for robustness.

Together, these resources form the foundation of a robust analytical framework designed to uncover clinically relevant drug repurposing opportunities and establish a scalable methodology applicable to other diseases. The approach centers on three key components: (1) extraction and preprocessing of genomic data, (2) analysis, identification, and validation of HNC-specific genotypic features, and (3) connecting with DrugBank to pinpoint viable drug repurposing opportunities. A clear diagram of the pipeline can be seen in **Figure** 1. A breakdown of the literature validation pipeline is available in **Figure 2**a.

**Figure 1.**
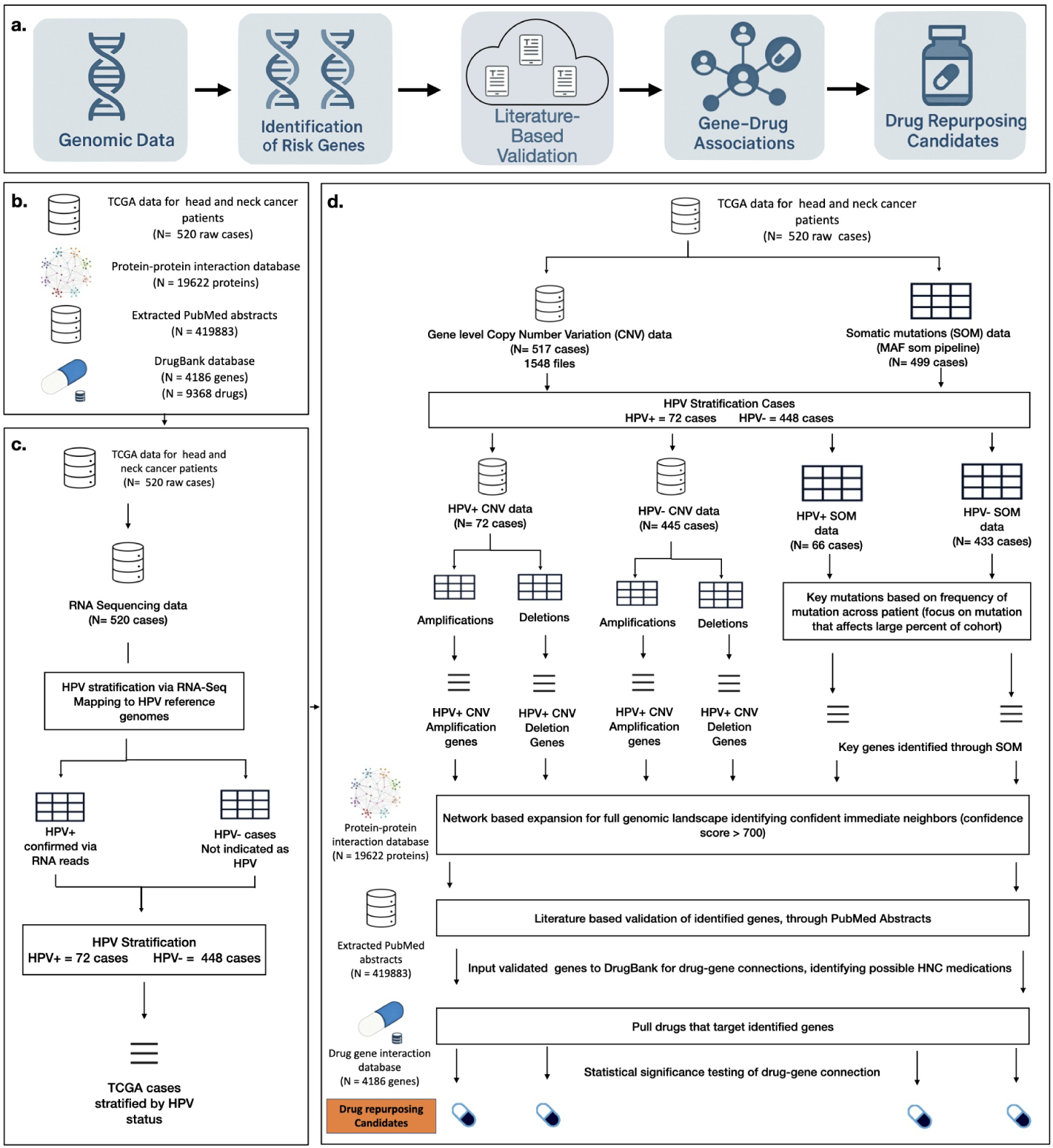
Overview of the GARD analytical pipeline for identifying drug repurposing candidates in HNC. **(a)** Illustrates the overall workflow, which begins with acquiring key genomic data, followed by identifying high-confidence genes, validating genes through literature-based analysis, and mapping them to drugs through curated gene–drug associations, ultimately yielding candidate medications for repurposing. Each section of this workflow is further broken down in the following section of the diagram which begins with **(b) input data sources**, which include multi-omics data from TCGA HNC cohort (520 raw cases curated from Case ids from Nulton et al [13]), the STRING PPI network comprising approximately 19,622 human proteins, extracted PubMed abstracts totaling 419,883 raw abstracts associated with HNC, and the DrugBank drug–gene interaction database containing 9,368 drugs and 4,186 protein targets [12,25,29]. Next, **(c) HPV stratification** is based on work completed by Nulton et al [13]. In which RNA-Seq data was used by extracting unmapped reads and aligning them to a reference panel of 171 HPV genomes, confirming HPV positivity primarily for HPV16, the predominant genotype in HNC, and classifying cases into HPV-positive (n = 72) and HPV-negative (n = 448) subgroups. Finally, **(d) the drug repurposing pipeline** integrates genomic alteration analysis, where copy number variation (CNV) and somatic mutation (SOM) data are processed to identify significant amplifications, deletions, and mutations across HPV-stratified cohorts. High-confidence risk genes are determined through statistical significance testing using binomial and permutation-based approaches with false discovery rate correction, then expanded through high-confidence PPI networks (STRING score 700) to capture pathway-level relevance. The genes were then validated through a literature-based validation step, leveraging publicly available sources (e.g., PubMed) to confirm drug–gene associations using recent research evidence. These validated gene sets are cross-referenced with DrugBank to identify drugs targeting either direct risk genes or their immediate network neighbors, and candidate drugs are determined using enrichment-based statistical tests, including hypergeometric and permutation analyses, ensuring significant drug-gene connections and resulting in a set of both approved therapies and novel repurposing opportunities.

**Figure 2.**
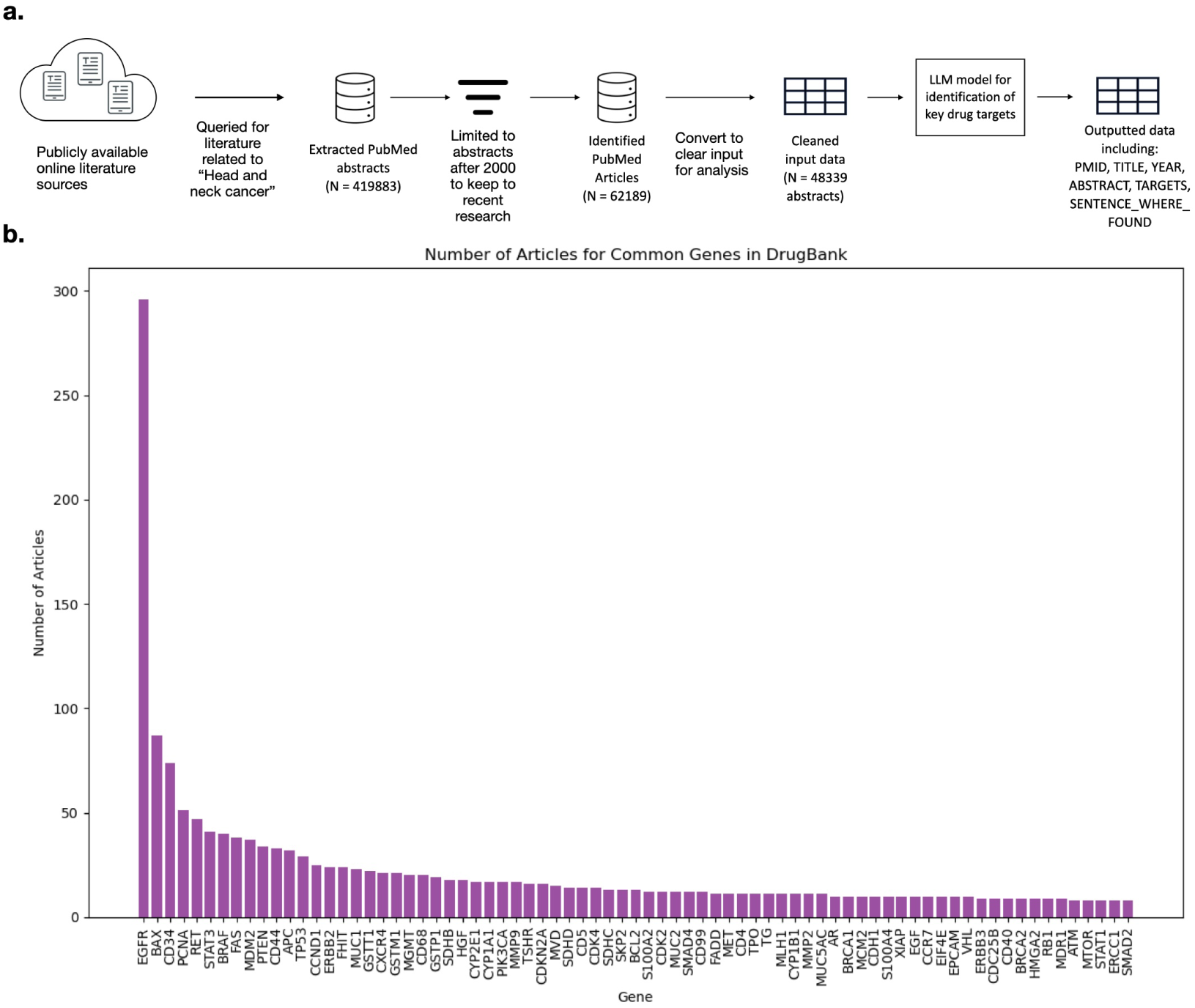
Literature-Based Validation Pipeline and Identified Gene Targets for HNC. **(a)** The validation pipeline workflow. The diagram illustrates a multi-step workflow designed to complement genomic analyses by leveraging published research. The process begins with querying PubMed for large-scale acquisition of biomedical abstracts related to HNC using NCBI’s E-utilities, followed by filtering to include only recent literature (published after the year 2000). The curated data is then formatted into structured tables to optimize interpretation by a large language model (LLM). Using Google GEMMA, abstracts are parsed to extract key therapeutic targets and supporting evidence. The final output integrates gene targets, source references, and associated drugs. **(b)** Top 75 gene targets ranked by literature support. Genes are displayed in descending order based on the number of PubMed articles referencing their association with HNC. Highly studied targets such as EGFR, TP53, PIK3CA, and CDKN2A dominate the top ranks, while emerging candidates with fewer publications highlight novel therapeutic opportunities.

### 2.2. TCGA data

The key genomic data was sourced from TCGA, a comprehensive and collaborative initiative between the National Cancer Institute (NCI) and the National Human Genome Research Institute (NHGRI): https://www.cancer.gov/tcga [12]. For this study the TCGA HNC cohort corresponding to case IDs from the Nulton et al study was utilized, allowing for HPV stratification. The final cohort included 520 cases, which were further refined based on the availability within each data category: copy number variation (CNV), and somatic mutation (SOM) data [12,13]. All data was accessed through the Genomic Data Commons (GDC) Data Portal. This study utilized open-access data, which includes de-identified clinical information, gene-level copy number variation (CNV) and somatic mutation profiles (SOM).

### 2.3. HPV stratification

HPV serves as an influential factor in the diagnosis and treatment of HNC [1,2,30, 31]. Its importance within the GARD pipeline was no different. HPV stratification was performed using results from Nulton et al., who demonstrated that RNA-Seq data can be used effectively to determine HPV status in HNC [13]. The majority of HPV-positive cases were HPV16, which was the primary genotype analyzed due to its prevalence and clinical relevance. Other HPV types were detected with low frequency. The Nulton et al. study analyzed 520 TCGA HNC cases and identified 72 HPV-positive patients [13].

### 2.4. Genomic Amplifications and Deletions

After stratification on HPV status, it was possible to utilize the data to identify key genomic components of HNC. TCGA provides gene-level copy number variation (CNV) data, which quantifies changes in the number of copies of specific genes across tumor samples. Deviations from this baseline, either deletions (CNV <2) or amplifications (CNV>2), can indicate genomic alterations that are possibly associated with the cancer or disease being studied.

Based on the categorization from Nulton et al. the key genetic mutations were categorized into HPV+ and HPV-cohorts. Within the extracted HNC TCGA cohort about 517 cases had cleaned CNV data. Within the available data for each sample, 58,645 available genes were analyzed and provided chromosome number, start and end position, and copy number value [12,32,33].

After identifying a gene as being amplified or being deleted it was important to determine whether the mutation was random or of significant prevalence across the entire disease landscape. In order to first filter to significantly altered regions for HNC a significance test for high level amplifications and deletions was calculated. High level amplifications were considered as those with a CNV value greater than 4. Along the same lines high level deletions were considered as those equaling 0 [34]. To determine significance of the mutation the counts of high-level amplification or deletion were utilized, to assess whether their occurrence across samples exceeded what would be expected by chance.

A binomial test was conducted where the significant mutation count per gene is the number of success across all files. This approach enabled derivation of a *p*-value to assess whether the CNV frequency in each gene exceeded what would be expected by chance given the cohort’s overall genomic instability. The calculation of this value was:

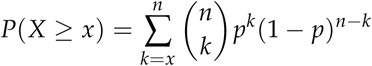

where *n* is the total number of samples in the cohort, *x* is the number of samples with high-level CNV amplification (CNV copy number *>* 4) or deletion(CNV *<* 1) events, and *p* is the genome-wide background CNV rate calculated as 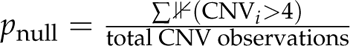 for amplifications, and 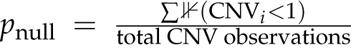 for deletions. This represented the proportion of all CNV measurements across all genes and samples that exceed the high-level threshold. This approach tested each gene against the observed cohort-specific genomic instability.

Alongside this an empirical null distribution derived through permutation testing. For each of 1,000 permutations (selected to balance computational burden and significance), *n* CNV values were randomly sampled (with replacement) from the pooled set of CNV observations, and the count of sampled values exceeding the threshold (CNV > 4, CNV<1) was recorded to build a null distribution. The empirical *p*-value was calculated by comparing the observed count to this null distribution. This was then divided by the number of permutations of the null distribution (n = 1000) to derive an empirical *p*-value. The empirical *p*-value was calculated as:

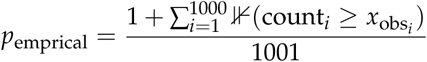

where *x*_obs_ is the observed number of samples with high-level CNV events for the gene, count*_i_* is the simulated count of high-level events in permutation *i*, ⊮ is an indicator function that equals 1 when the simulated count equals or exceeds the observed count, and the “+1” in both numerator and denominator prevents *p*-values of exactly zero while maintaining proper probability bounds. The two *p*-value calculations were conducted across many genes, and each test had its own chance of producing false positives. As such a false discovery rate (FDR) adjustment was necessary. The Benjamini-Hochberg correction was applied for this study[35].

To further filter key mutations a methodology was developed based on the methodology within the GISTIC software [34]. The key aspect is that mutations are assigned a score that considers both amplitude and frequency. Amplification values accumulated as log2 of the CNV value minus 1 to adjust for the diploid baseline. This helped to center the data around 2 or a diploid CNV count. A cap was placed on CNV values (CNV = 7) to ensure that outliers do not skew values and reduce visibility of genes with lower CNV amplitude but higher frequency. Deletion values were accumulated as +1 for minimal deletions (CNV=1) and +2 for significant deletions (CNV = 0) to track the amplitude of CNV alteration. The frequency of significant mutations was calculated by tracking the number of times a gene was significantly amplified (CNV >4) or deleted (CNV< 1). For amplification calculations the score was calculated as the average log2-adjusted amplification value (amplitude) multiplied by the frequency percentage (prevalence) of high-level mutations. For deletions the GISTIC-like score was calculated as the average absolute deletion intensity (amplitude) multiplied by the frequency percentage (prevalence) of significant deletion (CNV =0)[34].

Significant genes were first filtered to those with significant mutations according to the FDR adjusted *p*-values in both the binomial and empirical significance tests. To further refine candidate selection, the distributions of GISTIC like scores for these genes were evaluated to assess the strength of copy number alterations. Upper cutoff thresholds were determined directly from the observed score distributions, ensuring that only top CNV mutations were retained. By integrating statistical significance with genomic alteration burden, this process prioritized a focused set of high-confidence genes most relevant to HNC. These key genes could serve as potential targets for medications and in turn identify possible repurposing candidates [5,36].

### 2.5. Somatic Mutations

In addition to the copy number variations (CNV), somatic mutation (SOM) data was utilized. Somatic mutations were identified using cohort-level MAF analysis files from TCGA [13]. This identified cohort included information on a total of 16,477 mutations for the 499 cleaned cases available within the data. This data was stratified by the HPV status as described above, leading to 66 HPV positive cases and 433 HPV negative cases. After stratifying the data by HPV status, key genes were tracked by monitoring their mutations, using a method similar to the CNV tracking approach described in the above section.

The mutations were filtered to focus on non-synonymous mutation types as synonymous mutations rarely impact protein function [37]. Gene lengths were extracted from GENCODE v48 annotations and used to normalize mutation counts, as longer genes have greater opportunity to accumulate mutations by random chance simply due to larger mutational target size[38]. Each gene in the available data was analyzed to count the number of times a mutation occurs to a gene, as well as the unique number of patients harboring a mutation for the gene. Producing a mutation count and cohort frequency that were normalized by gene length.

A binomial test was then conducted to measure the enrichment of each mutation within the cohort and the significance of the mutation. Through the mutation based analysis, the mutational burden of each gene within the cohort could be directly measured. This test observed whether the mutation count in a gene significantly exceeds the expected count based on its genomic target size. The test assumes the observed mutation count as the number of successes and the total number of mutations across the cohort as the trial size. The test statistic was calculated as:

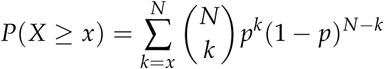

where *N* is the total number of mutations observed in the cohort (including both synonymous and non-synonymous mutations to establish an unbiased background rate),*x* is the observed number of non-synonymous mutations in the gene, and *p* is the gene-specific probability calculated as 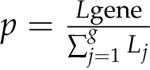 gene, where *L*_gene_is the coding sequence (CDS) length of the gene and the denominator represents the sum of CDS lengths across all protein-coding genes in the reference genome. This length-based analysis accounts for the fact that longer genes present larger mutational target spaces and are expected to accumulate more mutations by chance. The test directly measures whether the mutational burden in each gene exceeds expectations given its genomic target size.

As performed in the above section, an empirical significance test was conducted as well. For each gene a multinomial null distribution is constructed based on the mutation cohort, probability of gene mutation based on gene length calculated as gene length/sum of gene lengths. The null was constructed for each gene to be the size of the number of simulations (n = 10,000). A multinomial distribution was used to account for different mutation probabilities based on gene lengths.

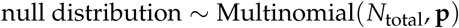

where *N_total_* is the total number of mutations observed across the cohort and **p** is the gene-specific probabilities, with 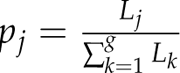 based on the CDS length *L_j_* of each gene. The multinomial distribution enables simultaneous simulation of mutation counts across all genes while preserving the constraint that total mutations remain constant and respecting differential mutational target sizes. The empirical *p*-value was calculated as:

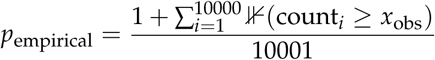

where *x*_obs_ is the observed mutation count in the gene and ⊮ is an indicator function that equals 1 when the simulated count equals or exceeds the observed count. As before, both binomial and empirical *p*-values were FDR corrected using the Benjamini–Hochberg procedure. Genes with FDR adjusted *p*-values < 0.05 in both tests were retained as significantly mutated, ensuring stringent control over false discoveries while maintaining adequate statistical power [35]. These statistical tests were utilized to remove background noise and filter to significantly mutated genes in the studied cohorts. From here, candidates were identified using similar methodologies to CNV analysis. A composite mutation score was calculated for each gene as:

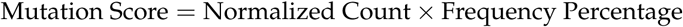

where Normalized Count represents the mutation burden per gene adjusted for coding sequence length 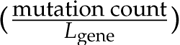, and Frequency Percentage is the proportion of patients harboring mutations in the gene. This scoring framework, balances mutation amplitude (burden per gene) with cohort prevalence (frequency across patients), ensuring that both recurrent and high-impact mutations are captured.

The distributions of normalized cohort frequency, normalized gene mutation count, and mutation score were evaluated to establish upper cutoff thresholds. These thresholds were determined directly from observed score distributions to identify genes with both statistical significance and substantial biological impact. Through integration of statistical significance with genomic alteration burden, this process prioritized a focused set of high-confidence genes that could be used to identify treatments for HNC and possible repurposing candidates.

### 2.6. STRING protein-protein interaction (PPI) network

Through the above methods, it was possible to identify key drug targets directly related to HNC. However, beyond the primary key genes directly implicated in HNC, their immediate network neighbors, genes connected either upstream or downstream in the relevant biological pathways, were also included in the analysis. This approach allowed for the consideration of both identified HNC direct risk genes and those with a single-degree interaction in the pathway context. Identifying these neighbors helps to account for other key genes that could potentially serve as drug targets [5,39]. Within the GARD pipeline, the STRING PPI network was utilized [25]. For this study the STRING database was filtered to focus on homo sapiens (human) gene interactions and pathways.

The direct risk genes identified through CNV and SOM mutations analysis were inputted to the STRING PPI to identify immediate neighbors of the genes. Once immediate neighbors were identified for the key genomic mutations the top connections were limited based on their confidence score. Based on the STRING db documentation a cutoff of 700 was used to keep immediate neighbors with high confidence PPIs [25]. The culminated top connected genes and key mutations of HNC were then utilized with the drug gene interaction database to identify medications that interact with these key targets.

### 2.7. LLM-based literature extraction for gene validation

The literature-based validation pipeline was designed to complement genomic analyses by leveraging published research. The validation data set was derived from publicly available resources in PubMed, a comprehensive repository including peer-reviewed biomedical and life sciences literature. The process begins with the large-scale acquisition of PubMed biomedical abstracts using NCBI’s E-utilities, a suite of tools designed for searching, retrieving, and analyzing PubMed records [27,40]. A query of “Head and Neck cancer targets” was provided to PubMed to extract all relevant literature for HNC key target identification. The query returned 419,883 abstracts, including titles, publication years, and abstracts. The dataset was then cleaned to remove incomplete or missing abstracts and filtered to include only studies published within the past 25 years (i.e., after the year 2000), ensuring relevance and contemporary scientific context. After cleaning and filtering, a total of 48,339 unique abstracts were retained for analysis.

The data was then structured for input to the model which included essential metadata such as PMID, title, publication year, and abstract text, as shown in **Table** 1. After compiling and formatting the data that included abstracts related to HNC the LLM Google GEMMA was utilized. Google GEMMA 2B instruct version was utilized. Google GEMMA has proven ability to interpret text and extract key information [28,41]. Due to the computational intensity of processing such a large dataset, the University of North Carolina at Chapel Hill (UNC) Longleaf High-Performance Computing Cluster was utilized to ensure efficiency and scalability. The LLM was prompted to identify key therapeutic targets related to HNC and extract supporting evidence by locating the specific sentence within each abstract where the target was mentioned. The output structure, illustrated in **Table** 2 was designed to facilitate rapid manual review and verification of model predictions before validating gene disease associations. Each output entry included the identified gene target, supporting sentence, source PMID, and associated drugs, enabling transparent validation and downstream analysis. An overview of this design analysis can be seen in Figure 2a.

**Table 1.** Input structure for LLM-Based Literature Parsing. This table shows the standardized input format provided to Google GEMMA for parsing PubMed abstracts. Each record includes the PMID, article title, publication year, and abstract text, ensuring structured and interpretable data

| PMID | TITLE | YEAR | ABSTRACT |
| --- | --- | --- | --- |
| Pubmed ID associated with the paper that was used | Title of the paper | Year of Publication | Abstract of the paper covering an overview of the paper and its associated results |

**Table 2.** Example Output Structure from LLM-Based Literature Parsing. This table illustrates the standardized format used to present results from Google GEMMA’s parsing of PubMed abstracts. Each entry includes the identified gene target, the supporting sentence from the abstract, the source publication (PMID), and associated drugs interacting with the gene. This structured approach facilitates efficient manual validation and downstream integration into drug repurposing analyses.

| PMID | TITLE | YEAR | ABSTRACT | TARGETS | Sentence where found |
| --- | --- | --- | --- | --- | --- |
| Pubmed ID associated with the paper that was used | Title of the paper | Year of Publication | Abstract of the paper covering an overview of the paper and its associated results | Targets identified as relevant targets for HNC through LLM analysis of Abstract | Key evidence in the abstract where the targets were found to be associated with HNC |

### 2.8. Drug-gene connection for drugs in DrugBank

After integrating immediate neighbors identified through high-confidence STRING PPIs and validating genes through literature-based analysis, both the top risk genes and their protein neighbors were queried against DrugBank to identify drug–gene associations [25,29]. The inclusion of PPI neighbors ensured that indirect but biologically relevant HNC genes were captured broadening the potential landscape of repurposable drugs. We then asked whether the HNC risk genes and their interacted protein neighbors are also drug target genes in DrugBank. If so, the statistical significance of drug–gene connections was reported. DrugBank is a curated database, providing detailed annotations on experimentally validated drug–gene interactions, including mechanism of action (e.g., inhibitor, activator, modulator). It contains information on 9,368 medications and 4,186 genes totaling 23,136 drug-gene interactions.

To assess the statistical significance of drug–gene connections, two complementary enrichment approaches were applied. First, a right-tailed hypergeometric test via cumulative binomial distribution was used to evaluate whether a given drug targeted more risk genes than expected by chance, given all druggable genes in DrugBank as the the background of the test. The method quantifies the likelihood of seeing the observed or more overlap in gene targets between the drug’s targets and risk genes, and to determine if this is more than expected by chance. Here, the null hypothesis stated that a drug’s target genes overlapped with the risk gene set randomly, while the alternative hypothesis indicated significant enrichment. The test statistic was calculated as:

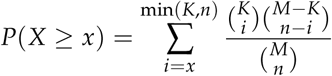

where *M* is the total number of druggable genes in DrugBank (4,186), *K* is the number of targets for a given drug, *n* is the number of risk genes (including PPI neighbors) present in DrugBank, and *x* is the observed overlap between the drug’s targets and the risk set.

Second, an empirical permutation test was conducted to strengthen confidence in the results. In this test, random gene sets of equal size to the risk set were repeatedly sampled (10,000 permutations) from the background universe of drug targetable genes (all available genes within DrugBank). For each permutation, the overlap with drug targets was calculated. The empirical *p*-value was then defined as the proportion of permutations with an overlap greater than or equal to the observed overlap. This distribution-based approach controlled for potential biases in gene–drug network structure. The empirical *p*-value was then defined as:

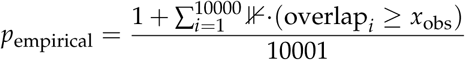

where *x*_obs_ is the observed overlap and the indicator function ⊮· equals 1 when the permuted overlap equals or exceeds the observed overlap. This distribution-free approach provides robustness against deviations from hypergeometric assumptions and accounts for heterogeneity in drug target set sizes.

Both hypergeometric and empirical *p*-values were FDR corrected for using the Ben-jamini–Hochberg procedure. Drugs with FDR adjusted *p*-values < 0.05 in both tests were retained as significant candidates [35]. This approach enabled the filtering of medications to identify those with significant drug-gene interactions. In conjunction with a set of genes validated through CNV and SOM analyses, as well as their immediate neighbors, these drugs demonstrate a strong association with HNC. The medications hold potential as therapeutic potential, those not originally indicated for HNC could serve as repurposing candidates.

## 3. Results

### 3.1. Overview of the GARD pipeline

We developed the GARD pipeline to perform a comprehensive analysis of HNC using genomic data from TCGA, stratified by human papillomavirus (HPV) status. Through GARD’s multi-step framework, integrating copy number variation (CNV), somatic mutation (SOM) profiles, PPI networks, and curated drug–gene associations from DrugBank, the pipeline identifies both established and novel genetic targets with therapeutic potential. An overview of the pipeline can be seen in Figure 1.

Genomic alterations were categorized into three major types: copy number amplifications, copy number deletions, and somatic mutations. Each category was analyzed independently to identify statistically significant gene-level changes associated with disease progression. High-confidence associated genes were identified using a combination of binomial and empirical permutation testing, with FDR adjusted *p*-values. This allowed for the removal of background noise and to filter to significantly mutated genes. Top significant genes were further filtered based on composite scores developed based on a combination of mutation amplitude (strength of mutation) and frequency (number of cases affected). These genes were then expanded through PPI networks in the STRING database using PPI combined score >700 as the cutoff for high-confidence interaction in PPI. This allows for the inclusion of immediate neighbors that may play biologically relevant roles in HNC pathogenesis [25].

After that, a key step within GARD of literature-based validation was performed for the above genes using PubMed abstracts parsed by the Google GEMMA LLM, which strengthens confidence in gene–disease associations and prioritizes biologically relevant targets [26,28]. Through large-scale analysis of PubMed abstracts, a comprehensive set of 6,077 unique genes associated with HNC was identified. Figure 2b illustrates the top 75 genes, sorted by article frequency, highlighting the most extensively studied targets in HNC. The identified genes include well-established oncogenic drivers such as EGFR, TP53, PIK3CA, and CDKN2A, which are frequently implicated in tumor progression and therapeutic resistance. Beyond these well-characterized genes, the analysis also uncovered novel or underexplored targets with limited prior association to HNC, offering opportunities for innovative therapeutic strategies. These findings confirm the robustness of the pipeline in capturing both key cancer genes and less-studied targets with potential biological significance. This literature-based validation step strengthens confidence in the genomic findings by providing independent evidence from peer-reviewed research.

Validated genes were then mapped to drug–gene associations using DrugBank, identifying both direct drug targets and network-derived candidates across HPV-positive and HPV-negative cohorts. To assess the statistical significance of drug–gene connections, two complementary enrichment approaches were applied: a right-tailed hypergeometric test to evaluate whether a given drug targeted more risk genes than expected by chance, and an empirical permutation test randomly sampling genes from the DrugBank universe compared observed overlaps with risk genes to that of the null distribution. This integrative approach revealed distinct molecular vulnerabilities within each subgroup and prioritized medications with strong mechanistic relevance to the identified gene sets. The following sections incorporate these methods to identify results and repurposing candidates.

**Table** 3 summarizes the number of high-risk genes identified through CNV and somatic mutation analyses, their connected immediate neighbors, and the corresponding significant medications uncovered through enrichment-based filtering.

**Table 3.**
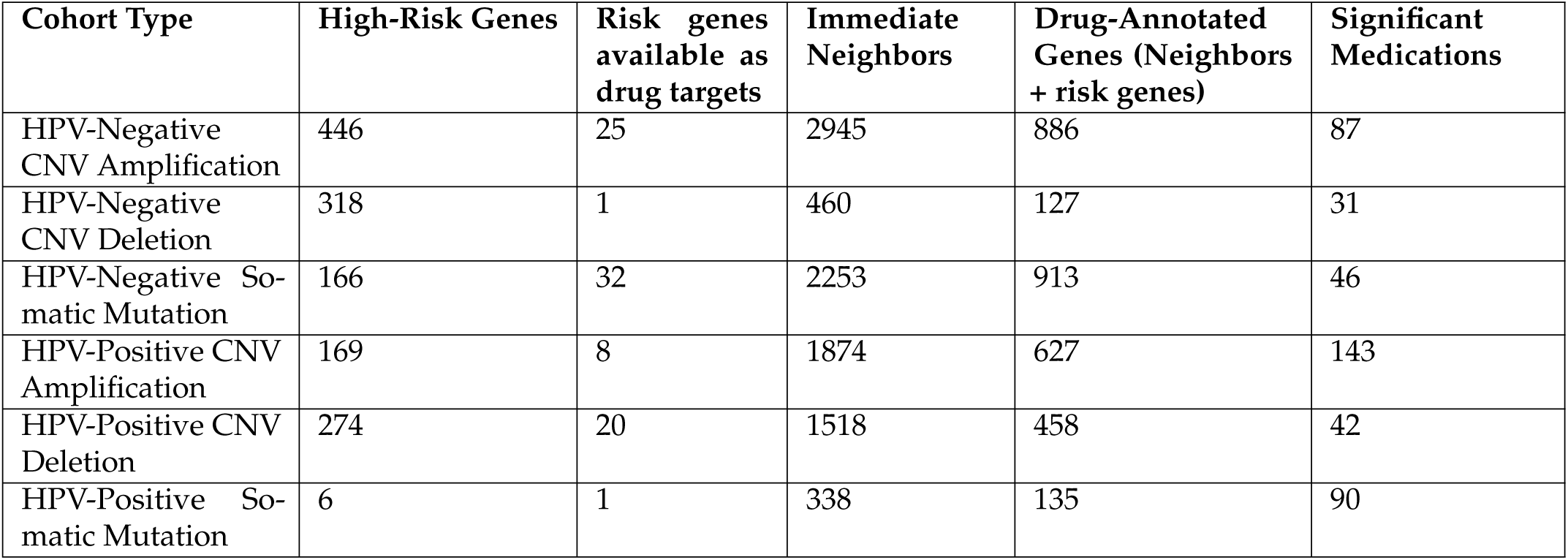
Summary of genomic and pharmacological analysis across HPV-stratified HNC cohorts. This table presents the number of high-risk genes identified through copy number amplification, deletion, and somatic mutation analyses in both HPV-positive and HPV-negative subgroups. It includes counts of direct drug targets, immediate neighbor genes identified via STRING PPI networks, drug-annotated genes from DrugBank, and the final number of statistically significant medications identified through enrichment-based filtering. These results provide a comparative overview of the molecular landscape and therapeutic potential across stratified HNC populations.

| Cohort Type | High-Risk Genes | Risk genes available as drug targets | Immediate Neighbors | Drug-Annotated Genes (Neighbors + risk genes) | Significant Medications |
| --- | --- | --- | --- | --- | --- |
| HPV-Negative CNV Amplification | 446 | 25 | 2945 | 886 | 87 |
| HPV-Negative CNV Deletion | 318 | 1 | 460 | 127 | 31 |
| HPV-Negative Somatic Mutation | 166 | 32 | 2253 | 913 | 46 |
| HPV-Positive CNV Amplification | 169 | 8 | 1874 | 627 | 143 |
| HPV-Positive CNV Deletion | 274 | 20 | 1518 | 458 | 42 |
| HPV-Positive Somatic Mutation | 6 | 1 | 338 | 135 | 90 |

### 3.2. HPV-Positive Investigation

Analysis of HPV-positive samples using CNV and somatic mutation data identified distinct genomic alterations associated with head and neck cancer. **Table 3** summarizes the cohort-level results, showing that the HPV-positive CNV amplification analysis yielded 169 high-risk genes, of which eight were available as drug targets, and led to 1,874 immediate neighbors totaling 627 targetable genes, resulting in 143 significant medications. The CNV deletion analysis identified 274 high-risk genes, of which 20 were available as drug targets, and led to 1,518 immediate neighbors totaling 458 targetable genes, resulting in 42 significant medications. Somatic mutation analysis revealed six high-risk genes, of which one was available as a drug target (PIK3CA) and led to 338 immediate protein neighbors totaling 135 targetable genes, resulting in 90 significant medications from DrugBank. Overall, combination of the identified significant genes and their protein neighbors gave 177 drugs to be repurposed, which is still relatively large.

To further refine this candidate set literature validation of the genes was applied. A total of five genes (appendix Table A1) were validated in literature among the above 51 unique risk genes found to be targetable or connected to targetable genes through PPI. Among the validated risk genes, somatic mutations identified PIK3CA; amplification analysis derived PIK3CA, RFC4, CLDN1 and SOX2; and deletions identified TLR7. From here the 177 drugs were reduced to 84 medications, 6 of which were found through directly targeting risk genes (appendix Table A2) and 84 medications were found through indirect targeting of literature validated protein neighbors (appendix Table A3).

Drugs directly targeting risk genes are illustrated in Figure 3a, highlighting key genes with existing therapeutic agents and potential candidates for drug repurposing (Table in appendix Table A2). Among direct targets, PIK3CA was associated with medications such as Wortmannin, XL765(Voxtalisib), Buparlisib, Copanlisib, and CH-5132799. With the latter three identified as inhibitors. Buparlisib and Copanlisib have progressed to initial trials in combination regimens [42–45], CH-5132799 has shown potent antitumor activity in preclinical models though it remains investigational [46], and XL765 shows promising PI3K inhibition [47,48]. Wortmannin despite its PI3K inhibition, is limited by poor selectivity and toxicity[49]. In contrast, TLR7, was linked to Golotimod, an immune-modulatory agent that completed Phase II trials for oral mucositis in HNC patients [50].

**Figure 3.**
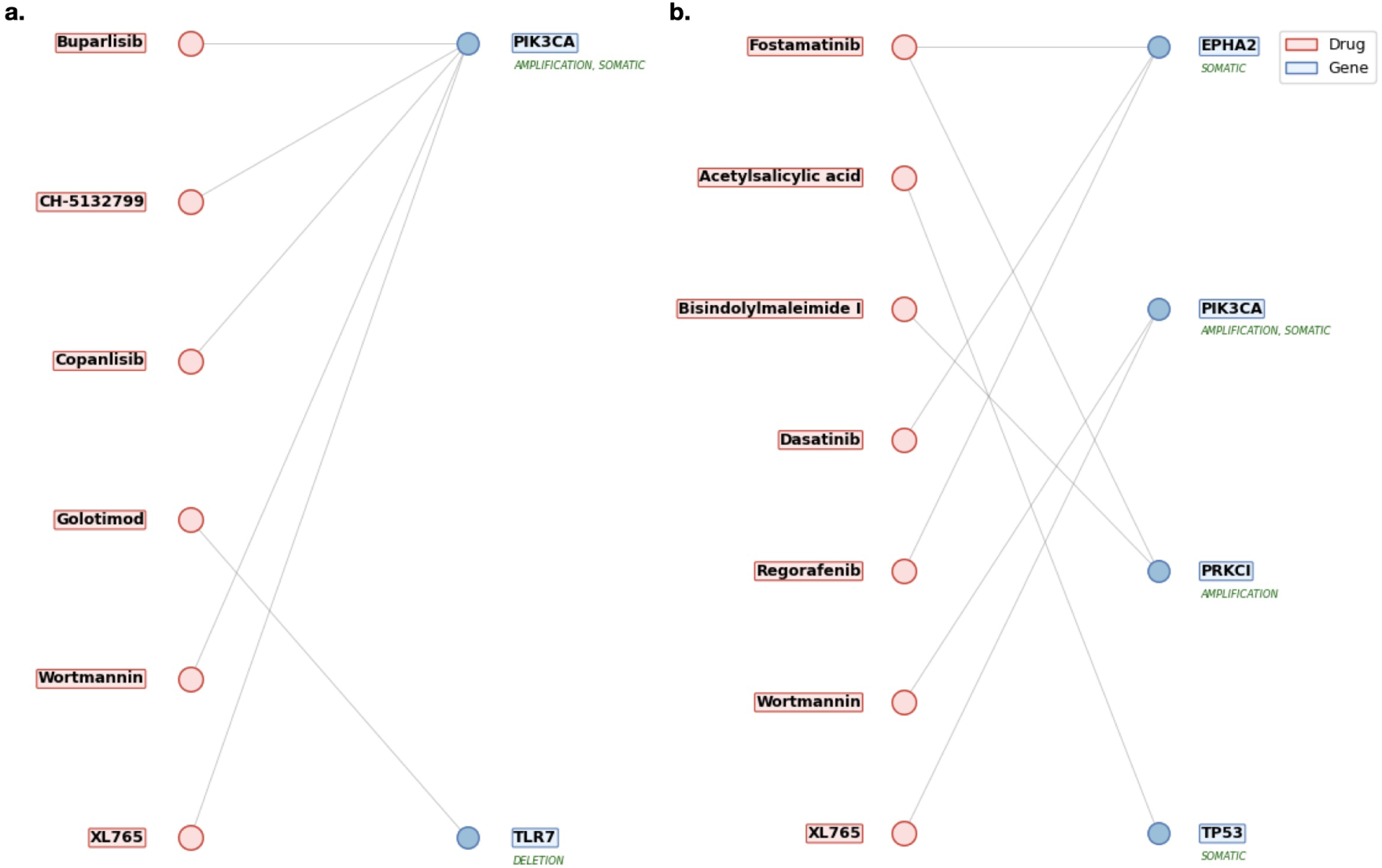
(a) Drug repurposing candidates targeting direct genomic risk genes in HPV-positive HNC. This figure presents medications identified through integrative analysis of copy number variations and somatic mutations in HPV-positive tumors. Each drug is linked to a direct genomic target, such as PIK3CA, or TLR7, that was statistically validated as a high-confidence driver of disease. The listed candidates represent potential therapeutic agents for precision oncology in this distinct HNC subgroup. Candidates included CH-5132799, Golotimod, Buparlisib, and Copanlisib with strong mechanistic ties to HNC treatment. **(b) Direct genomic targets identified in HPV-negative HNC from TCGA analysis.** This figure presents a curated list of medications identified through integrative genomic analysis of HPV-negative HNC tumors using TCGA data. It highlights drugs that directly target high-confidence genomic risk genes uncovered via copy number variation (CNV) and somatic mutation profiling. These candidates, including agents such as Fostamatinib and Regorafenib, demonstrate therapeutic relevance by interacting with key oncogenic drivers like TP53, PIK3CA, and EPHA2, offering promising avenues for precision drug repurposing in HPV-negative HNC. These linked to drug candidates of Acetylsalicylic acid, Regorafenib, Dasatinib, and Fostamatinib.

Furthermore, identified risk genes were expanded through high-confidence PPI networks, incorporating biologically relevant neighbors. Through validation of immediate neighbors and the direct risk genes they were connected to, the 263 unique druggable immediate neighbors originally identified were focused to 33. Figure 4 illustrates the relationships between drugs, their gene targets, and high-risk genes for top candidates identified by the pipeline. Top candidates were selected from the top 50 medications ranked by FDR adjusted significance of drug–gene associations in HPV-positive HNC (full list in appendix Table A3). The medications target critical oncogenic pathways stemming from connections to risk genes of PIK3CA, SOX2, and CLDN1 underscoring their role in HNC [51–54]. Notable candidates included kinase inhibitors such as Afatinib, Fostamatinib, Brigatinib, Cabozantinib, Capivasertib, Deuruxolitinib, Amuvatinib, Regorafenib, and Nintedanib that work to target critical nodes including EGFR, MET, RET, KIT, PDGFRA, IGF1R, ALK, JAK2/3, AKT iso-forms, and the ERBB, FGFR, NTRK families.

**Figure 4.**
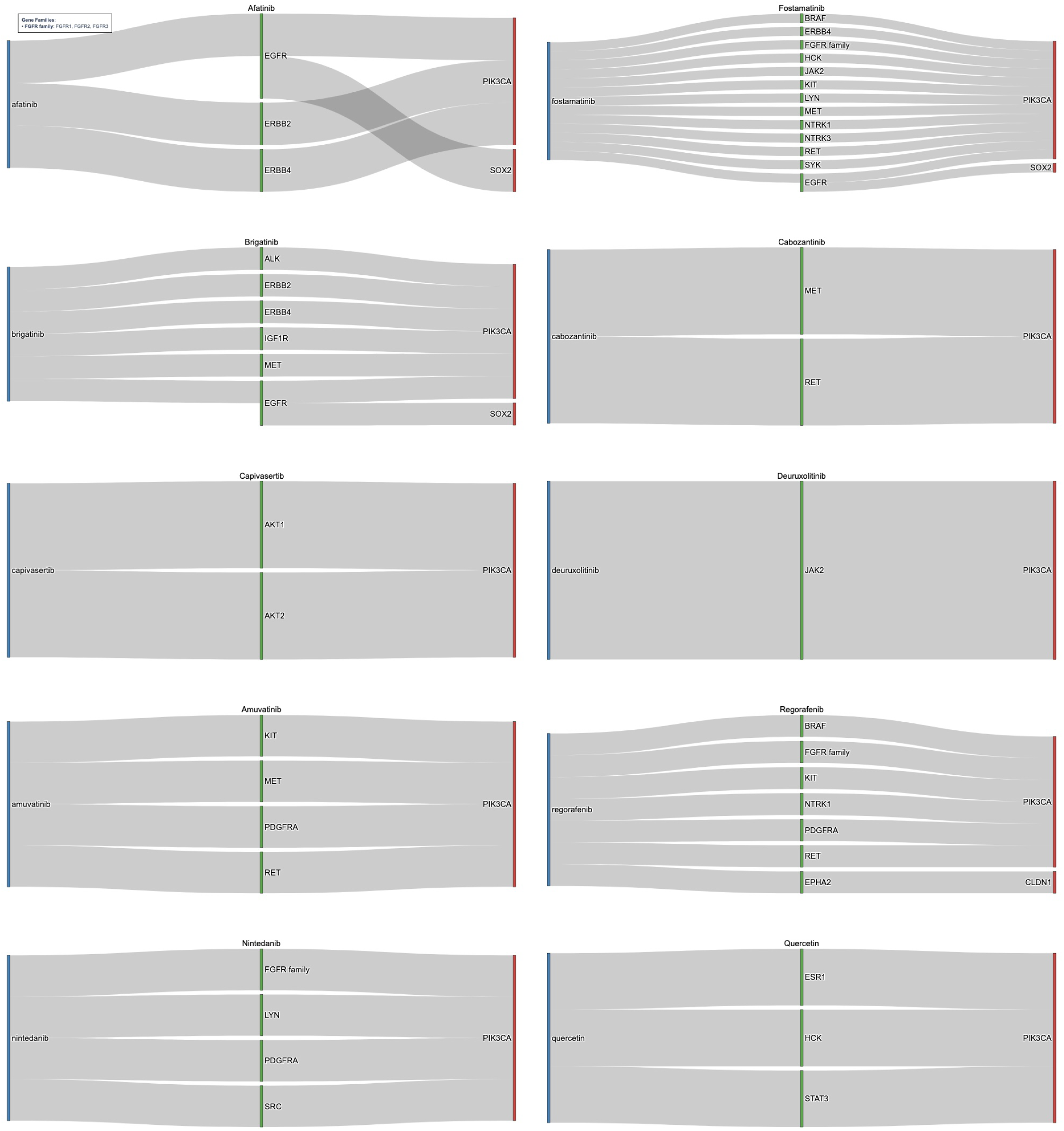
Sankey diagram for drug repurposing candidates target indirect genomic risk genes in HPV-positive HNC. This figure shows medications identified through network-based expansion of direct genomic targets using high-confidence PPI data. Each diagram shows the drug (blue) its gene target (green) and the risk gene (red), that was identified through CNV or SOM analysis, that is a neighbor to the gene target and led to its identification. The drugs target immediate neighbor genes, such as EGFR, STAT3, and the FGFR family, which are functionally connected to core drivers of HPV-positive HNC and represent biologically relevant therapeutic opportunities. These were found through connections to key risk genes of PIK3CA, SOX2, and CLDN1. In this manner kinase inhibitors such as Amuvatinib and Regorafenib alongside medications such as Quercetin were identified.

Afatinib, an EGFR/ERBB inhibitor, showed efficacy in recurrent/metastatic HNC in trials such as LUX-Head & Neck [55,56], while Cabozantinib improved progression-free survival when combined with cetuximab or immune checkpoint inhibitors[57–59]. Fostamatinib stands out for its broad multi-kinase inhibition and Phase II evaluation in solid tumors, including HNC [60]. Capivasertib, a potent AKT inhibitor, selectively targets AKT1 and AKT2 and has demonstrated preclinical efficacy in HNSCC models, highlighting its potential to disrupt PI3K/AKT-driven oncogenesis [61–63]. Deuruxolitinib, a JAK2 inhibitor, represents an emerging candidate due to its ability to modulate the JAK/STAT pathway, which is frequently activated in HPV-positive tumors and associated with immune evasion and inflammation, making it a rational choice for repurposing [3,64]. Nintedanib has progressed to Phase II clinical trials in HNC owing to its FGFR-targeting activity, addressing a key vulnerability in HPV-positive disease [65]. Brigatinib, originally approved for non-small cell lung cancer, inhibits ALK and EGFR signaling, suggesting strong potential for overcoming resistance mechanisms in HPV-positive HNC [66,67]. Amuvatinib, a multi-targeted tyrosine kinase inhibitor, acts on KIT, PDGFRA, MET, and RET, positioning it as a promising candidate for repurposing [68]. Regorafenib, interacts with KIT, PDGFRA, RET, EPHA2, and FGFR family members connected to PIK3CA and CLDN1, providing a mechanistic basis for its application in HNC [69]. Additionally, non-kinase agents like Quercetin, identified for its pro-apoptotic and antioxidant effects, found to target ESR1, HCK, and STAT3, linked to PIK3CA shows potenital for HNC treatment beyond conventional kinase inhibition [65].

Overall, HPV-positive tumors identified key risk genes that were expanded to included key signaling and targetable pathways, supporting the identification of novel therapeutics.

Many were known or in clinical trials, but those that were not held potential as repurposing candidates. These findings highlight the potential for rapid translation of repurposed drugs into clinical applications for HPV-positive HNC.

### 3.3. HPV-Negative Investigation

Similar to HPV positive analysis CNV and SOM analysis identified HNC associated genes. Table 3 shows that HPV-negative CNV amplification analysis identified 446 high-risk genes, of which 25 were available as drug targets, and led to 2,945 immediate neighbors totaling 886 targetable genes, resulting in 90 significant medications. CNV deletion analysis yielded 318 high-risk genes, of which one was available as a drug target, and led to 460 immediate neighbors totaling 127 targetable genes, resulting in 30 significant medications. Somatic mutation analysis revealed 166 high-risk genes, of which 32 were available as drug targets, and led to 2,253 immediate neighbors totaling 913 targetable genes, resulting in 42 significant medications. This led to a total of 117 unique medications to serve as repurposing candidates.

Through literature analysis this was further refined. A total of 20 genes (appendix Table A4) were validated among the 224 unique risk genes found to be targetable or connected to targetable genes through PPI. Among these genes somatic mutations identified TP53, PDE3A, MAGEC1, EPHA2, COL1A2, HRAS, REG1A, LRP1B, RAC1, CDKN2A, and PIK3CA. Amplifications derived CLDN1, DVL3, EIF4G1, PDCD10, PRKCI, RFC4, SOX2, and PIK3CA (identified through two methods). Deletions identified CDKN2B and CDKN2A (identified through two methods). This allowed the 117 medications identified to be reduced to 58 medications, 8 of which were found through directly targeting risk genes (appendix Table A5) and 58 were found through indirect targeting of validated protein neighbors (appendix Table A6).

Drugs directly targeting risk genes are shown in in Figure 3b, with full details pro-vided in appendix Table A5. Among the validated genes, EPHA2, PIK3CA, PRKCI, and TP53, were found to have significant relationships with medications. The medications directly targeting risk genes included some overlap with HPV+ direct risk gene targeting candidates such as Wortmannin and XL765, but included further candidates such as Fostamatinib, Acetylsalicylic acid, Bisindolylmaleimide I, Dasatinib, and Regorafenib. Although Wortmannin shows significant relationship to the risk genes it remains limited by poor selectivity and toxicity[49]. XL765’s appearance in both cohorts shows strong relationship to HNC related genes such as PIK3CA and potential for repurposing [47,70]. Fostamatinib remains a strong candidate as it has a broad applicability to many genetic targets related to HNC, it is currently in trials for HNC [60]. Acetylsalicylic acid commonly known as Aspirin is a pain reducer with anti inflammatory properties. With limited information in regards to HNC, initial findings prove promising and it has shown effectiveness in other types of cancers[71]. Bisindolylmaleimide I is a compound utilized for protein kinase inhibition. It is not currently used clinically, but warrants further investigation due to its interaction with PRKCI, but due to its lack of use as a medication remains as lower priority repurposing candidate [72]. Dasatinib is a tyrosine kinase inhibitor used in treatment of leukemia, and has reached phase II clinical trials for HNC [73]. Finally, Regorafenib appeared again and shows strong inhibition of EPHA2 [69].

Identified risk genes were expanded through PPI networks to incorporate relevant neighbors. Validation of immediate neighbors and the direct risk genes they were connected to reduced the 472 unique druggable immediate neighbors originally identified to a focused 77. Figure 5, which highlights the top medications identified after network expansion, shows the relationship between drugs, their gene targets, and root high-risk genes. These medications were selected from the top 50 medications ranked by FDR adjusted significance of drug–gene associations in HPV-negative HNC (full list in appendix Table A6). The top medications targeted pathways stemming from connections to risk genes of TP53, PIK3CA, EIF4G1, SOX2, HRAS, RFC4, RAC1, EPHA2, and CDKN2A. Notable candidates included multi-kinase inhibitors such as Fostamatinib, Regorafenib, Brigatinib, Erdafitinib, Lucitanib, Lenvatinib, and Nintedanib targeting validated neighbor genes like EGFR, SRC, KIT, RET, MET, HSPA8, STAT3, PDGFRA, and the FGFR, ERBB, and RPS families. Several of these agents, Fostamatnib, Regorafenib, Brigatinib, Nintedanib, were common to both HPV-positive and HPV-negative cohorts, reinforcing their broad mechanistic relevance. Others, such as Lenvatinib and Erdafitinib, emerged as novel candidates for HPV-negative disease. Lenvatinib, targeting the FGFR family, PDGFRA KIT, and RET, has demonstrated efficacy in HNC as part of combination regimens in clinical trials [74,75]. Erdafitinib, found to targets similar genes, shows potential for repurposing in solid tumors, including HNC [76]. Lucitanib was found to target the FGFR family and PDGFRA and has shown promise in clinical trials for HNC [77]. The candidates of Fostamatnib, Regorafenib, Brigatinib, and Nintedanib were found again between the HPV+ and HPV-cohorts and shown to interact with similar key genes such as FGFR family, KIT, RET, PDGFRA, and SRC [60,66,69,78, 79]. Beyond kinase inhibitors other significant medications were identified. Artenimol, originally developed as an antimalarial, exhibits anticancer activity through immune modulation, targeting genes connected to TP53 and EIF4G1 [80,81]. Quercetin, previously highlighted in HPV-positive analysis, reappeared as a candidate due to its role in PI3K/AKT and STAT3 signaling [65]. Finally, Acetylsalicylic Acid (Aspirin) was found to target pathways connected to TP53, HRAS, and PIK3CA, reinforcing its potential anticancer activity anticancer effects and candidacy for repurposing in HNC [71].

**Figure 5.**
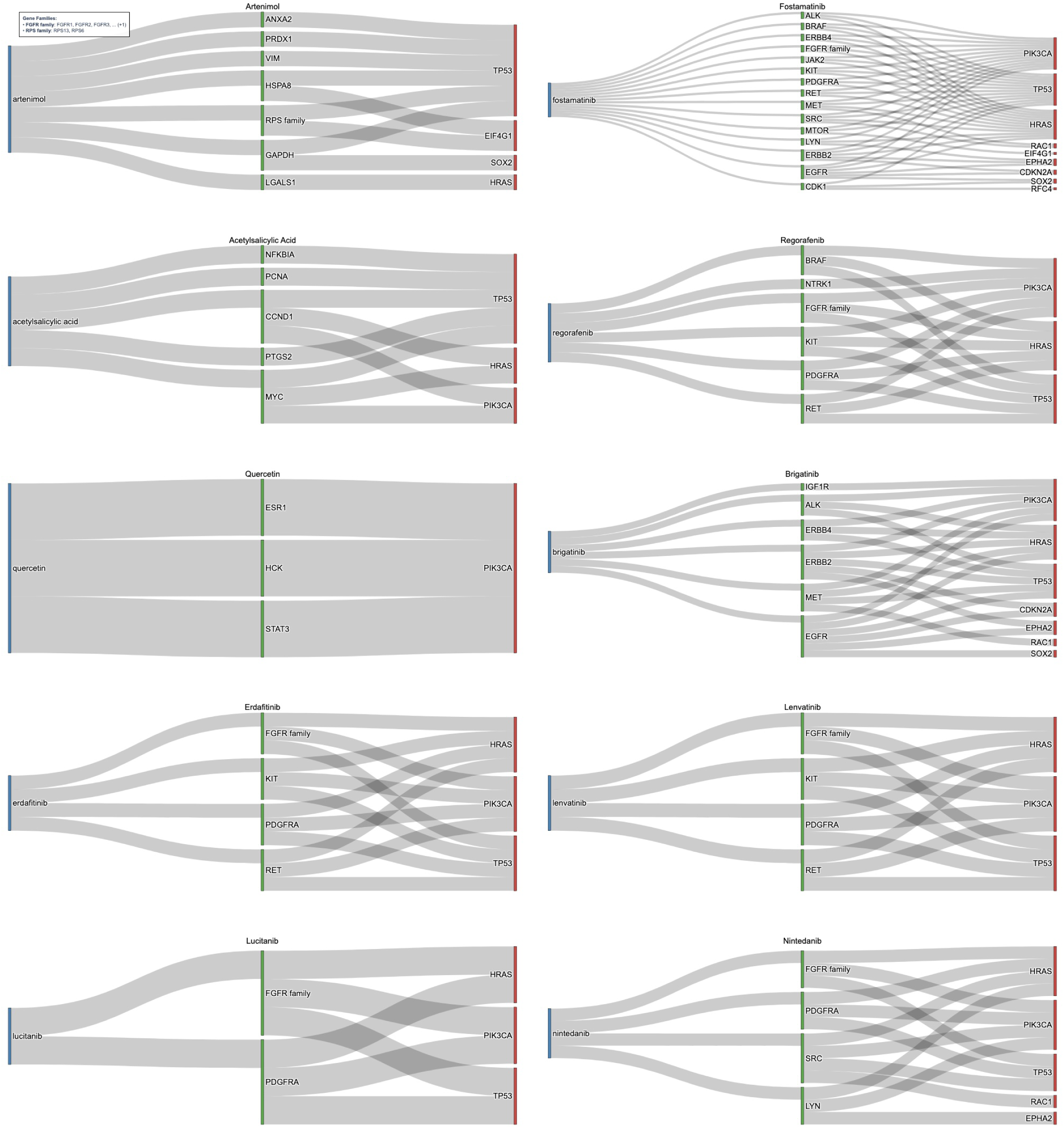
Sankey Diagram for Drug Repurposing Candidates Targeting Indirect Genomic Risk Genes in HPV-Negative HNC. This figure illustrates medications identified through network-based expansion of direct genomic risk genes in HPV-negative HNC using high-confidence PPI data from the STRING database. Each drug (blue) is connected to a gene target (green) that serves as an immediate neighbor to a risk gene, identified through CNV or SOM analysis, (red). The drugs target immediate neighbor genes functionally linked to core oncogenic drivers such as TP53, CDKN2A, and PIK3CA, representing biologically relevant therapeutic opportunities. Key candidates include multi-kinase inhibitors such as Lenvatinib and Erdafitinib, as well as drugs like Quercetin and Acetysalicylic Acid, which collectively offer promising strategies for precision oncology.

Through the identified risk genes alongside PPI expanded gene pathways key therapeutic targets were identified within the HPV-negative tumors. This led to the identification of known, or under investigation, medications alongside possible novel or exploratory repurposing candidates.

## 4. Discussion

With the expansion of available databases to utilize, drug repurposing has become a more popular and feasible way of discovering novel treatments for cancers and diseases [5]. This study introduces a genomics-driven, network-informed pipeline, GARD (Genomic Alteration-based Repurposing for Drugs), designed to systematically identify repurposing candidates for head and neck cancer (HNC). By integrating multi-omics data from HPV-stratified TCGA cohorts with high-confidence PPI networks and literature-based validation, GARD captures subtype-specific molecular signatures and expands them into biologically relevant networks. This multi-layered approach enables the identification of both direct genomic drivers and compensatory signaling hubs, which are then mapped to curated drug–gene associations from DrugBank, providing a robust framework for precision drug repurposing in HNC.

The analysis of the HPV-positive HNC cohort revealed a distinct molecular profile characterized by PIK3CA, SOX2, RFC4, CLDN1, and TLR7. Many were known cancer or HNC associated genes but incorporation of genes such as TLR7 presented underexplored targeting avenues [53,54,70,82–85]. These genes were expanded through PPI networks to further identify biologically relevant neighbors, and potential new therapeutic targets. These expanded genes included critical signaling mediators and pathway regulators (e.g., EGFR, KIT, JAK2/3, MET, RET, EPHA2, FGFR family, PDGFRA, and STAT3), which play essential roles in oncogenesis and therapeutic resistance [24,79,86–88]. Literature validation reinforced the mechanistic relevance of these targets, highlighting opportunities for precision drug repurposing in HPV-positive HNC. These results highlighted known pathways while uncovering novel targets, creating opportunities to identify repurposing candidates by either by leveraging newly discovered gene targets or by repositioning medications that act on established targets. The pipeline demonstrated its effectiveness by identifying drugs already in clinical trials for HNC, such as Buparlisib, Copanlisisb, Afatinib, Cabozantinib, Nintedanib, and Fostamatinib[42,45,56,59,78]. At the same time, it revealed promising new candidates, including XL765( Voxtalisib), Brigatinib, Amuvatinib, Deuruxolitinib, Capivasertib, and Regorafenib, which interact with critical signaling networks and represent opportunities for innovative repurposing strategies [47,64,66,68,69]. Beyond kinase inhibitors, Golotimod emerged as a unique candidate for immune modulation through TLR7 targeting [83,89]. Quercetin was also identified as a non-kinase agent with antioxidant and pro-apoptotic properties, offering unique potential to modulate PI3K/AKT and inflammatory pathways [65].

In HPV-negative tumors, alterations were dominated by cell-cycle regulators such as TP53 and CDKN2A/B, along with oncogenic drivers including PIK3CA, PRKCI, stem cell regulators like SOX2, tight-junction and EMT regulators such as CLDN1, and receptor tyrosine kinase nodes such as EPHA2 [14,53,54,70,86,90,91]. These genes were expanded through high-confidence PPI networks, revealing additional biologically relevant neighbors involved in critical pathways such as ERBB2/4, RET, KIT, EGFR, PDGFRA, MET, SRC, STAT3, FGFR family, and RPS family [24,79,87,88,92]. This network expansion broadened the therapeutic landscape by incorporating both known drivers and novel targeting opportunities not traditionally associated with HNC, such as EIF4G1. Literature-based validation reinforced the mechanistic relevance of these targets, highlighting opportunities for precision drug repurposing in HPV-negative disease. The pipeline confirmed its predictive strength by identifying drugs already in clinical trials for HNC, including Fostamatinib, Dasatinib, Lucitanib, and Lenvatinib [60,73–75,77]. At the same time, it uncovered novel candidates such as Brigatinib, Regorafenib, Erdafitinib, Amuvatinib, Artenimol, Acetylsalicylic acid, and Quercetin, which expand therapeutic options through possible repurposing candidates [47,64,66,68,69]. Notably, Erdafitinib, an FGFR inhibitor approved for bladder cancer, and Brigatinib, an ALK/EGFR inhibitor used in lung cancer, represent strong mechanistic candidates for repurposing in HPV-negative HNC. Beyond kinase inhibitors Quercetin emerged as an agent with antioxidant and pro-apoptotic properties, offering unique potential to modulate PI3K/AKT and inflammatory pathways [65]. Similarly, Artenimol, originally developed as an antimalarial, demonstrated anticancer activity through oxidative stress and immune modulation, while Acetylsalicylic Acid (Aspirin) offers a low-cost anti-inflammatory strategy with emerging evidence of anticancer benefits [71,80,81].

By integrating direct genomic targets with network-based expansion, the GARD pipeline broadens the identified genes beyond known drivers, incorporating biologically relevant neighbors. The inclusion of literature-based validation further strengthens confidence in these identified targets, reducing false positives and prioritizing high-confidence targets. Identifying these gene targets and protein neighbors associated with HNC provides insight into the disease enabling identification of drugs that can modulate these pathways, providing evidence for making them strong candidates for repurposing. This multi-layered approach allowed for the identification of known medications alongside novel candidates that have clear safety profiles and mechanistic pathways, improving the potential for translation into clinical practice. Importantly, the stratification of cohorts by HPV status proved essential for revealing distinct molecular vulnerabilities. HPV-positive and HPV-negative tumors exhibit fundamentally different drivers, which influence treatment response. By capturing these subtype-specific signatures, the GARD pipeline enables precision drug repurposing tailored to the unique biology of each subgroup, moving beyond one-size-fits-all approaches. Overall, the integration of multi-omics data, network-based expansion, and literature validation through the GARD pipeline provides a robust and scalable framework for systematic drug repurposing in HNC and broader application across other malignancies.

The study’s reliance on TCGA data introduces limitations related to cohort size and population diversity, as such expansion through new databases or larger cohorts could provide expanded results. Furthermore, while the computational predictions presented here are promising, experimental validation remains essential. Future work should focus on translating these predictions into preclinical models, such as HPV-stratified cell lines and adaptive clinical trial designs to confirm efficacy and their use in clinical practice as viable therapeutic options.

## 5. Conclusion

In conclusion, the GARD pipeline identified 84 candidate medications for HPV+ HNC and 58 medications for HPV-HNC. Many medications were in initial trials for HNC and showed viability of the pipeline for identification of repurposing candidates. Further underexplored medications have the potential to serve as repurposing candidates in the field of HNC. The GARD pipeline demonstrates the potential for genomic based drug repurposing and shows its potential in identifying novel treatments in HNC or further diseases.

## 6. Author Contributions

Pradham Tanikella (Conceptualization, Data curation, Formal analysis, Methodology, Validation, Software, Writing original draft, Writing review & editing), Will Nenad (Methodology, Writing review & editing), Christophe Courtine (Writing review & editing), Yifan Dai (Writing review & editing), Qingying Deng (Visualization, Writing review & editing), Baiming Zou (Writing review & editing), Nosayaba Osazuwa-Peters (Writing review & editing), Travis Schrank(Methodology, Writing review & editing), and Di Wu (Conceptualization, Formal analysis, Methodology, Validation, Supervision, Software, Writing original draft, Writing review & editing).

Conflict of interest: The authors declare no conflict of interest. The funders had no role in the design of the study; in the collection, analyses, or interpretation of data; in the writing of the manuscript; or in the decision to publish the results.

## Abbreviations

The following abbreviations are used in this manuscript: 
HNC: Head and neck cancer
TCGA: The Cancer Genome Atlas
HPV: Human Papillomavirus
PPI: Protein-Protein Interaction
FDR: False Discovery Rate
GARD: Genomic Alteration-based Repurposing for Drugs

## Appendix

This appendix provides a detailed extension of the main analysis by presenting comprehensive tabular outputs from the drug repurposing pipeline for HNC. It includes three key components for each HPV stratified cohort: (1) tables of top genes identified through copy number variation (CNV) and somatic mutation analyses, stratified by HPV status, which highlight statistically significant drivers of disease biology validated through binomial and empirical testing; (2) tables of direct drug–gene associations, listing medications that target these high-confidence genes, supported by literature evidence and prioritized using empirical FDR adjusted *p*-values; and (3) tables of indirect drug–gene associations derived from network-based expansion using high-confidence STRING PPI data, showcasing the top 50 medications ranked by empirical significance. Together, these resources provide transparency into the computational workflow, illustrate the molecular distinctions between HPV-positive and HPV-negative cohorts, and support the identification of both established and novel therapeutic candidates for precision oncology in HNC.

For the code utilized in this analysis pipeline please refer to the github link: https://github.com/pvtanike/Genomic-Landscape-Based-Drug-Repurposing.git

**Table A1.**
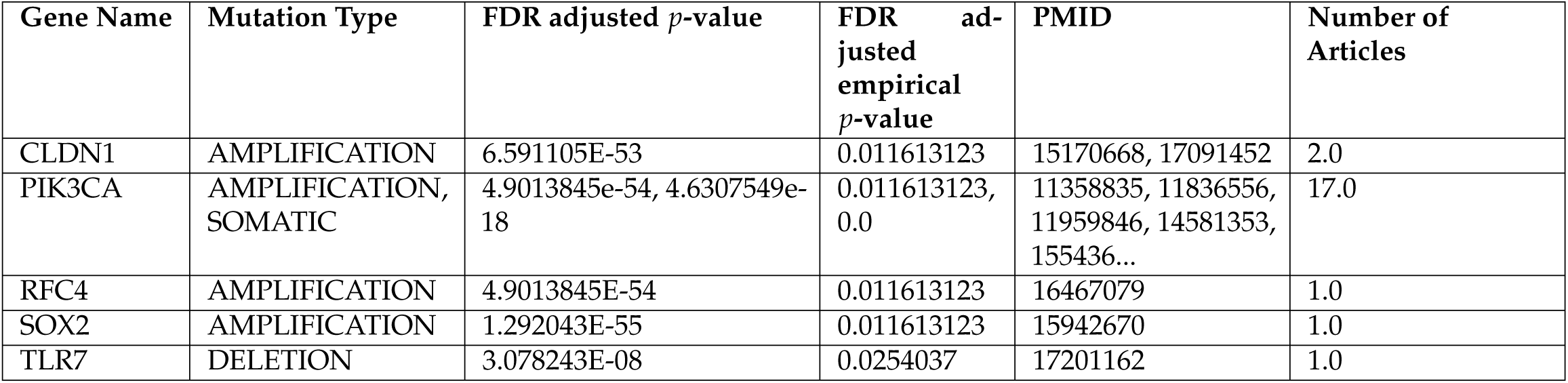
Top genes identified in HPV-positive HNC cohort, with columns for gene name, mutation type (MUT TYPE), q-value, and empirical q-value derived from permutation (CNV) or multinomial (SOM) null distributions. These genes represent statistically significant drivers of disease progression, identified through CNV and somatic mutation analyses with FDR adjustment. Their inclusion highlights core oncogenic and immune-related alterations, such as amplifications in PIK3CA and deletions in TLR7, that underpin the distinct molecular landscape of HPV-positive tumors. All listed gene targets were further validated through large-scale literature mining, ensuring strong evidence of their biological relevance in HNC. These findings informed downstream drug–gene mapping and network-based expansion, ultimately guiding the identification of precision drug repurposing candidates.

| Gene Name | Mutation Type | FDR adjusted *p*-value | FDR adjusted empirical *p*-value | PMID | Number of Articles |
| --- | --- | --- | --- | --- | --- |
| CLDN1 | AMPLIFICATION | 6.591105E-53 | 0.011613123 | 15170668, 17091452 | 2.0 |
| PIK3CA | AMPLIFICATION, SOMATIC | 4.9013845e-54, 4.6307549e-18 | 0.011613123, 0.0 | 11358835, 11836556, 11959846, 14581353, 155436... | 17.0 |
| RFC4 | AMPLIFICATION | 4.9013845E-54 | 0.011613123 | 16467079 | 1.0 |
| SOX2 | AMPLIFICATION | 1.292043E-55 | 0.011613123 | 15942670 | 1.0 |
| TLR7 | DELETION | 3.078243E-08 | 0.0254037 | 17201162 | 1.0 |

**Table A2.**
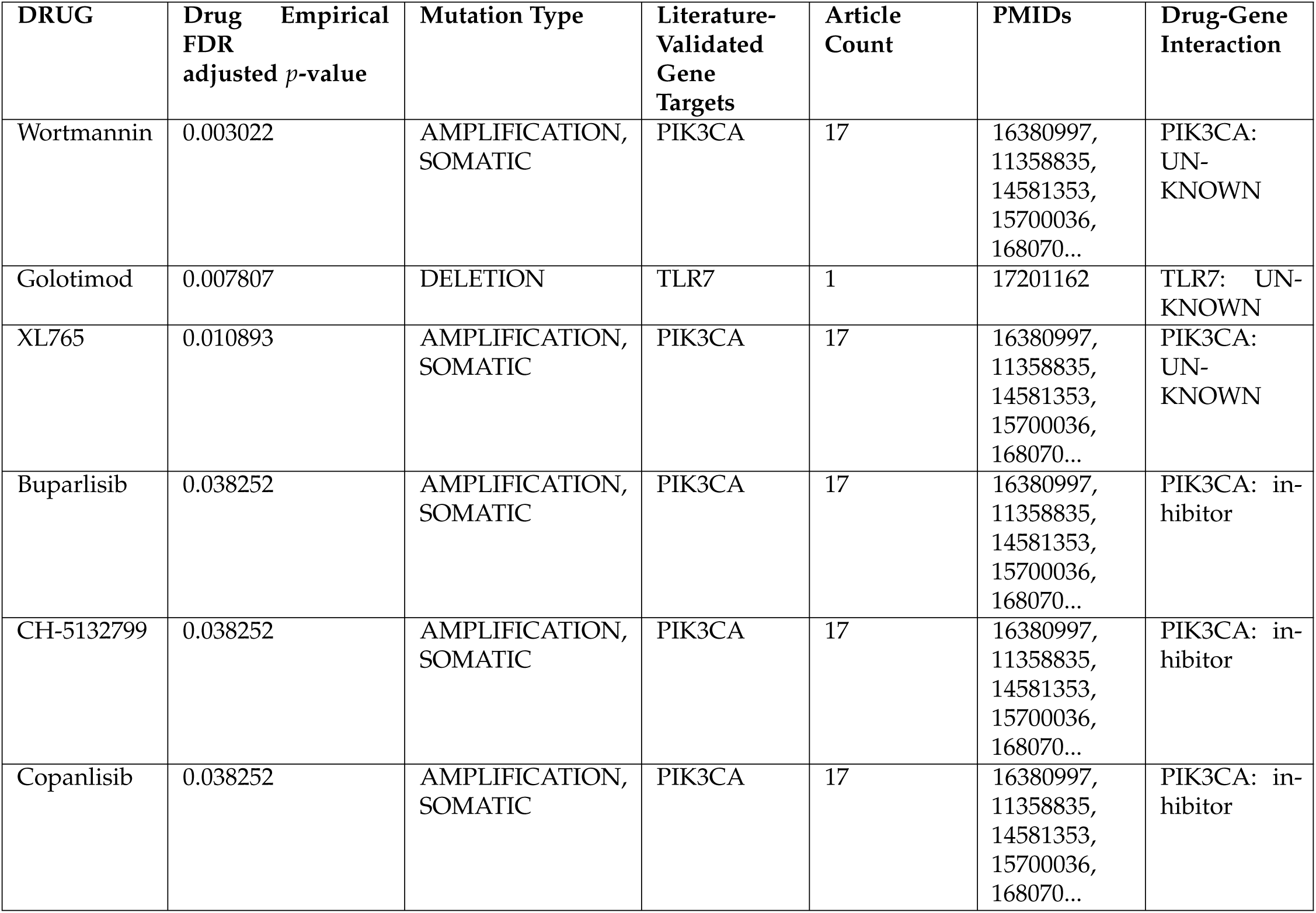
Direct drug–gene associations for the HPV-positive HNC cohort are summarized in this table. Columns include drug name, literature-validated gene targets, mutation type (MUT TYPE), supporting PMID references, number of articles, and empirical FDR adjusted *p*-values. These medications directly target high-confidence genomic alterations, such as amplifications in *PIK3CA* and deletions in *TLR7*, that define the molecular landscape of HPV-positive tumors. To ensure biological relevance, all risk genes were validated through a literature-based pipeline, and only candidates targeting genes with literature validated associations to HNC were retained. This validation step strengthens mechanistic confidence, while empirical FDR adjustment ensures statistical robustness. These direct associations informed the prioritization of top candidates, including PI3K inhibitors (Buparlisib, Copanlisib) and immune modulators (Golotimod), which emerged as promising repurposing opportunities for precision therapy in HPV-positive HNC.

**Table A3.**
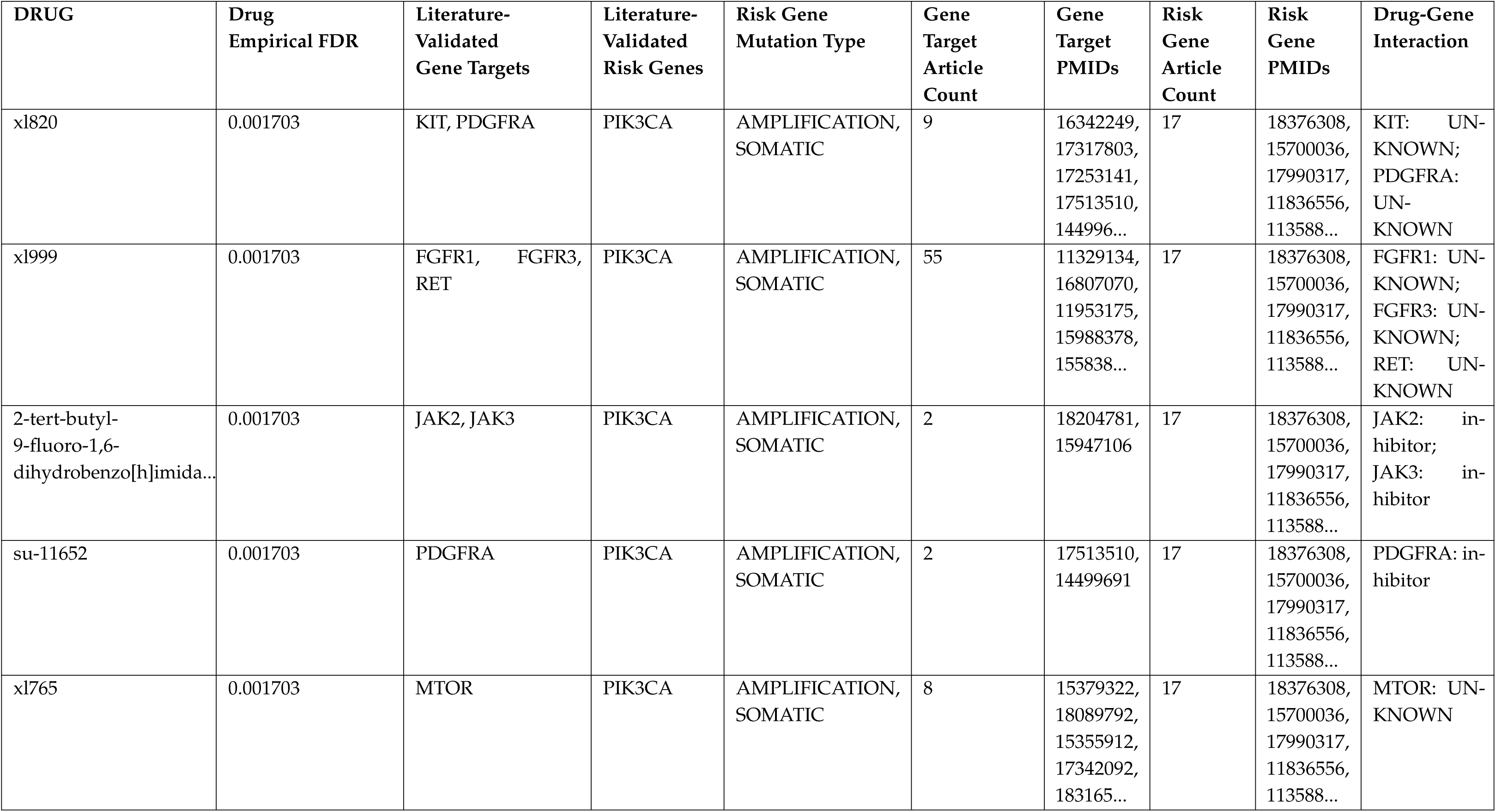
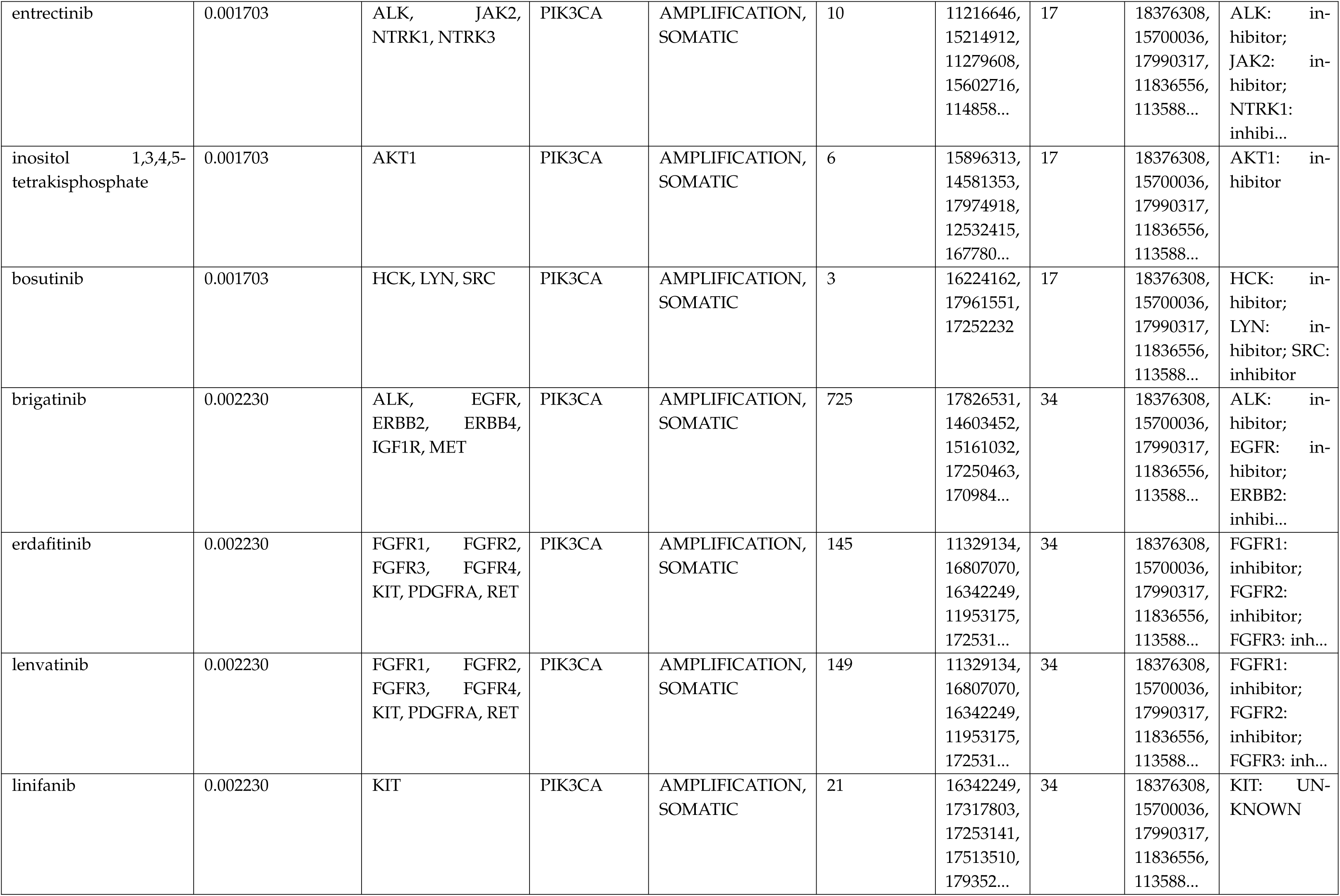
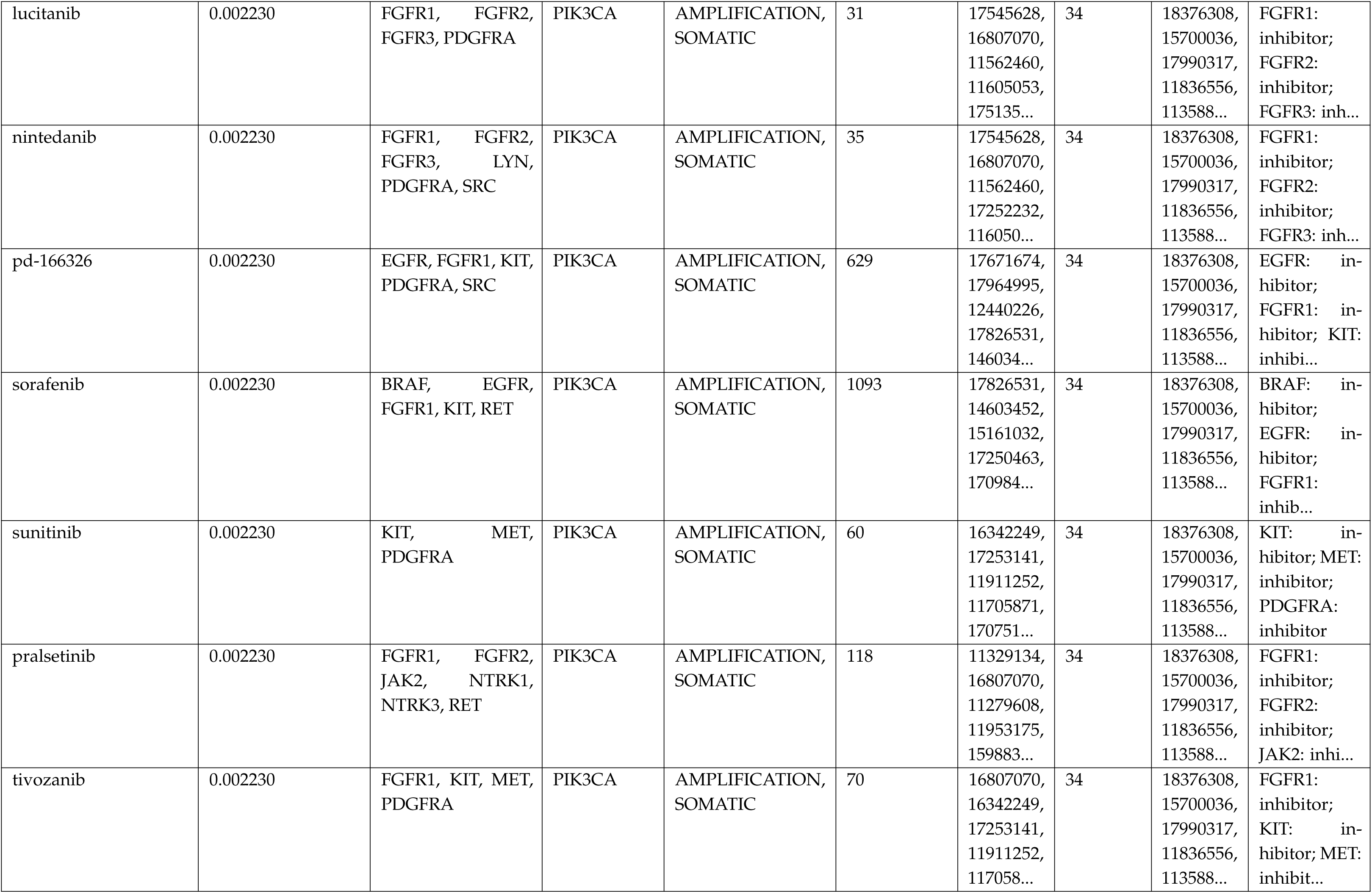
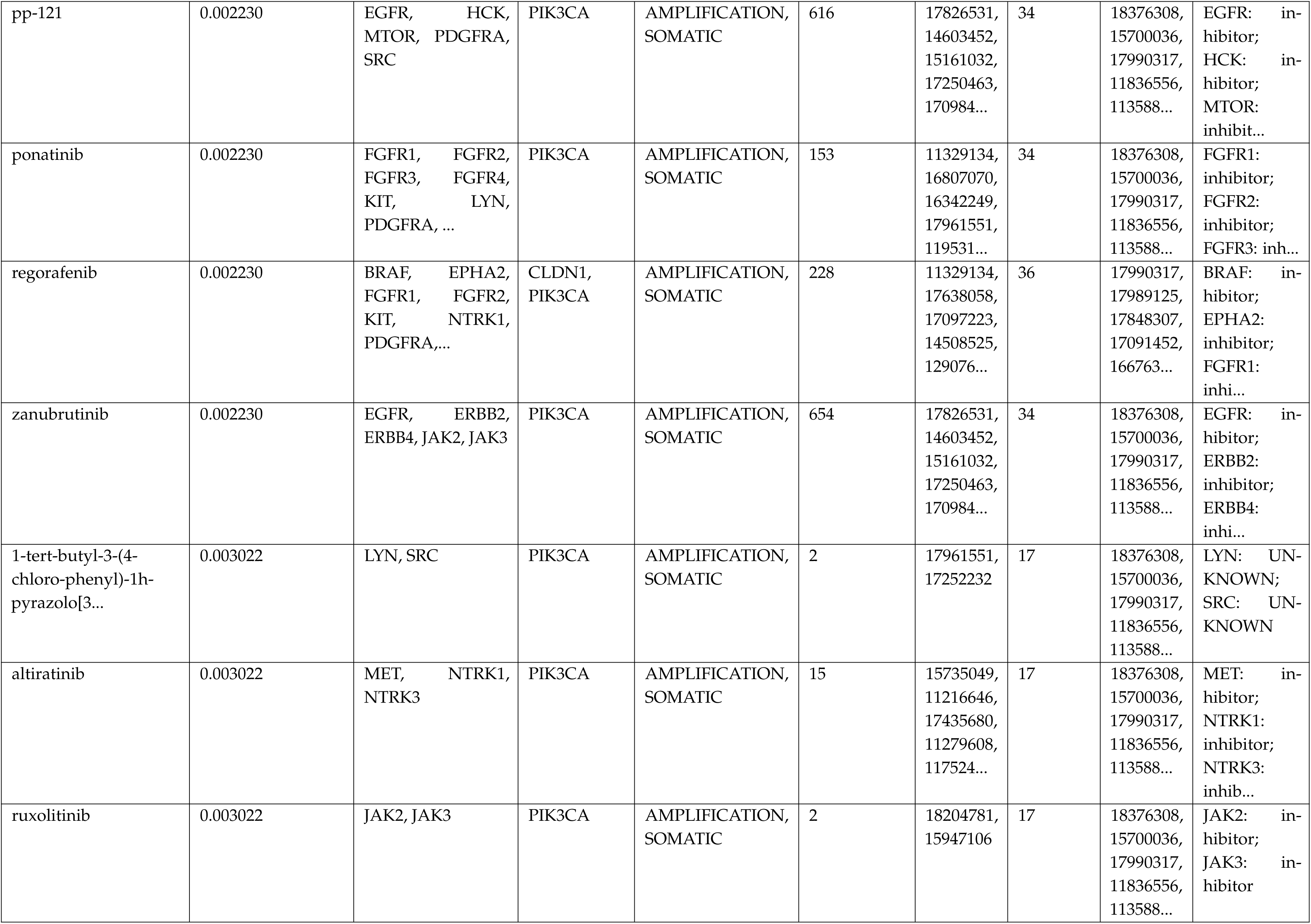
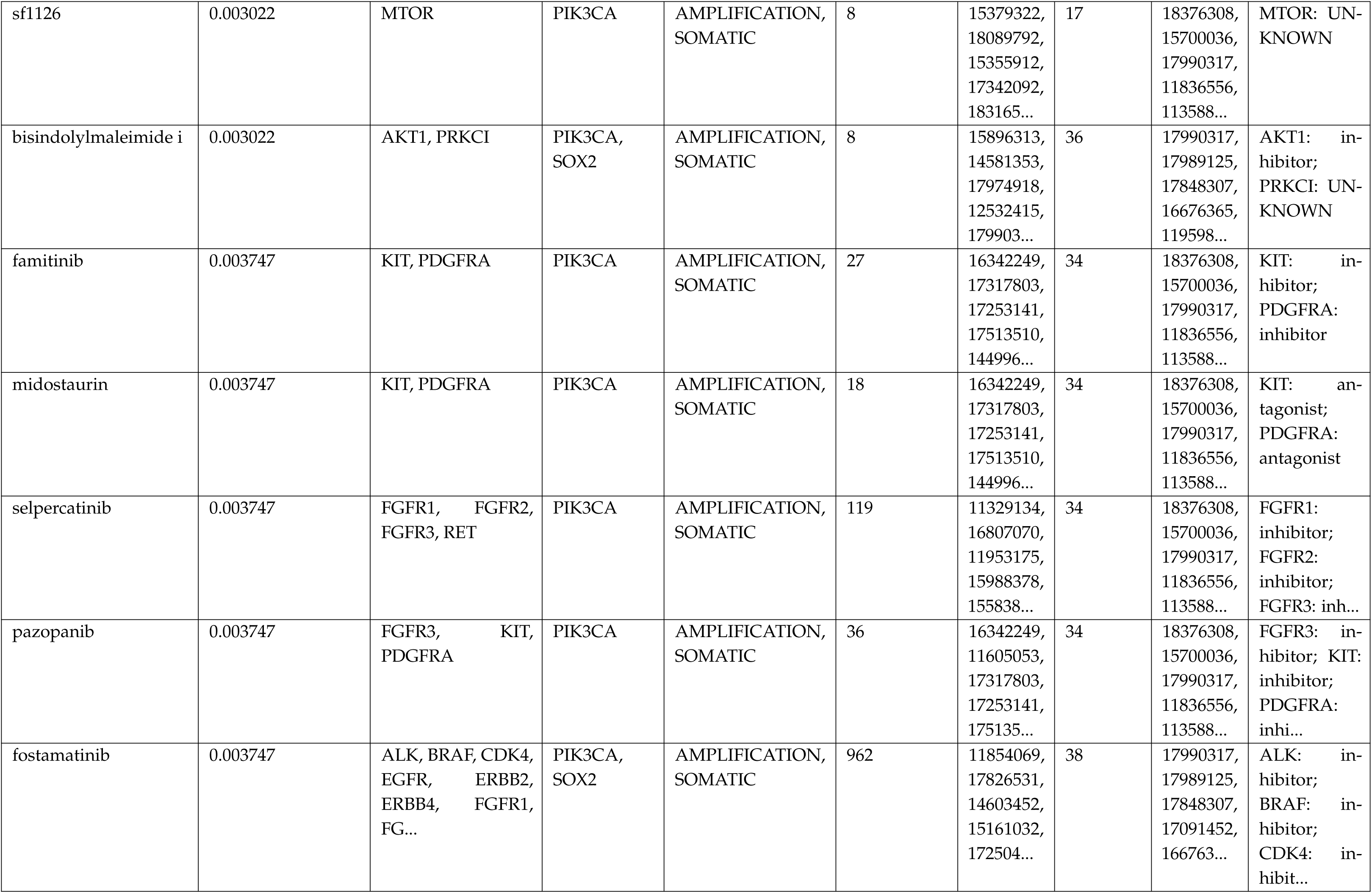
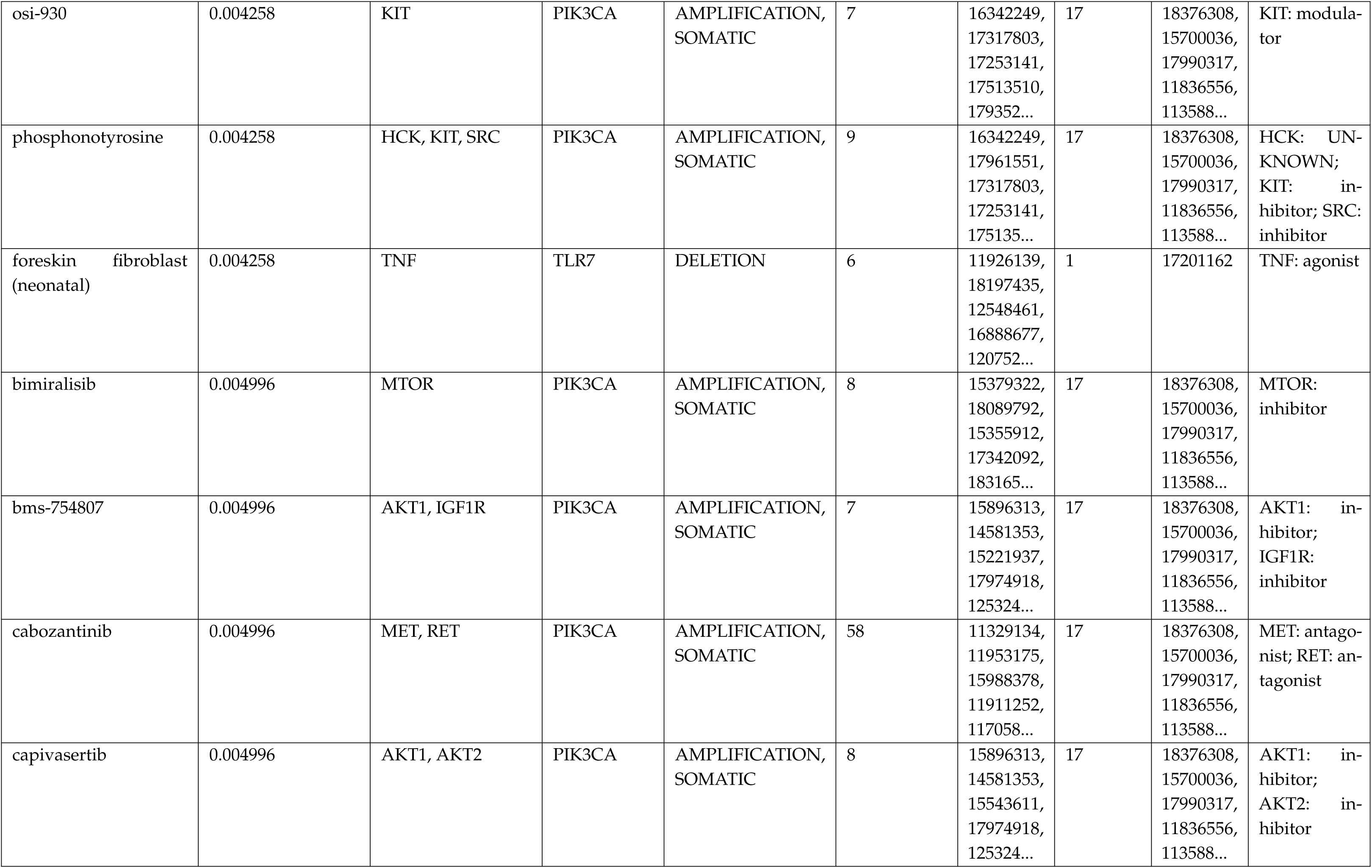
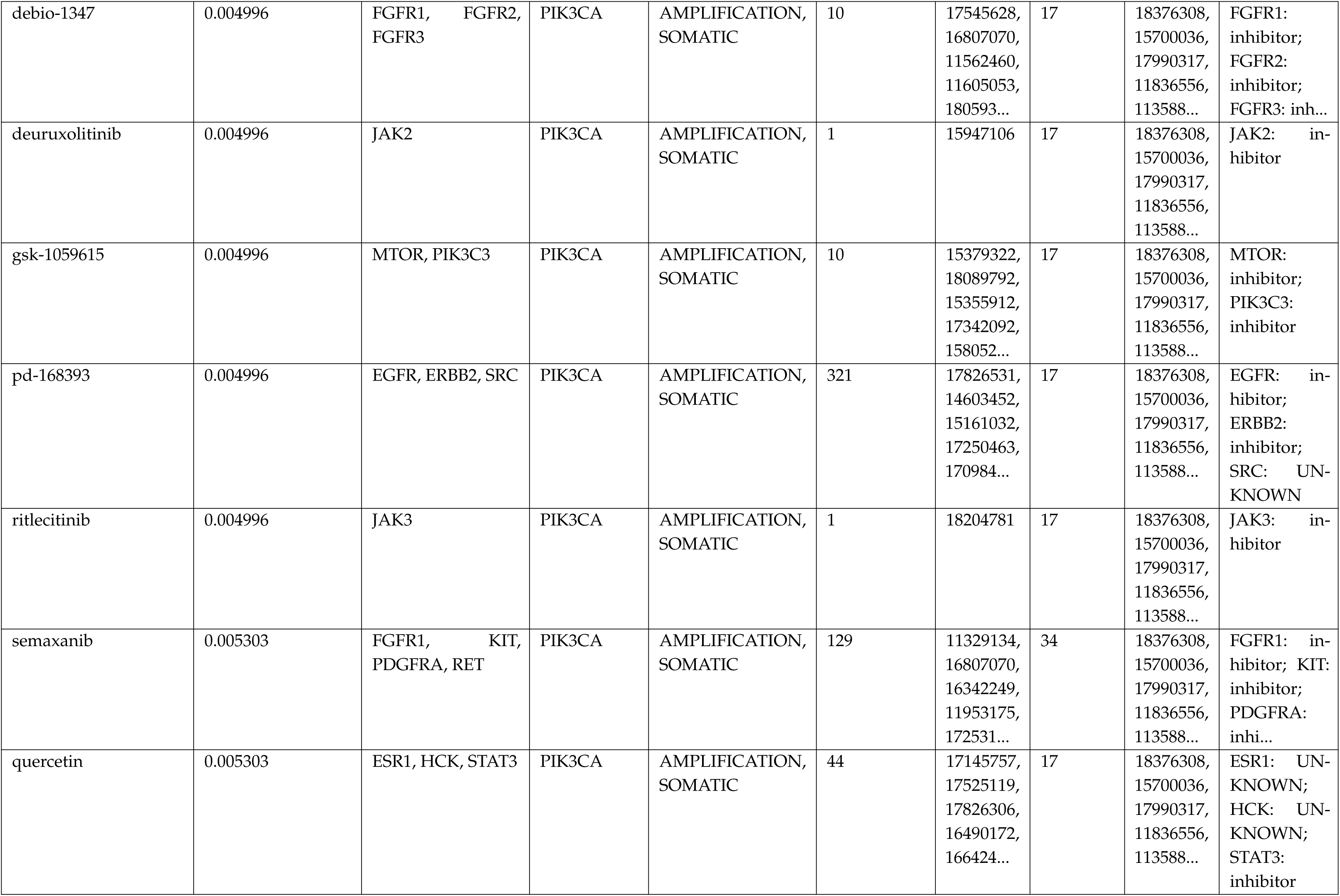
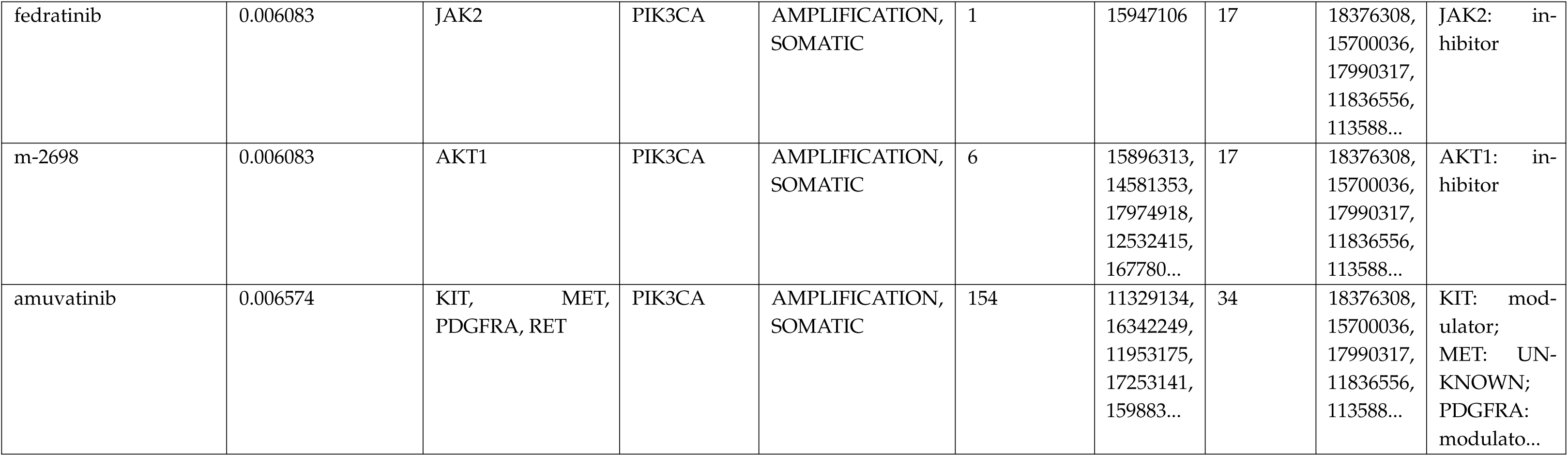
Top 50 indirect drug–gene associations for HPV-positive HNC cohort, ranked by empirical FDR-adjusted. *p***-values.** Columns include drug name, literature-validated gene targets, associated connected risk gene, mutation type (MUT TYPE), supporting PMID references, number of supporting articles, and empirical FDR-adjusted *p*-values. These medications target immediate network neighbors of high-confidence risk genes identified through CNV and somatic mutation analyses, leveraging STRING PPI data to capture pathway-level relevance. All gene targets and their network neighbors were validated through a large-scale literature mining pipeline to ensure biological significance, and only candidates with literature supported associations to HNC were retained. This network-based expansion broadens the therapeutic landscape beyond direct genomic alterations, uncovering multi-kinase inhibitors (e.g., Brigatinib, dacomitinib) and novel agents (Quercetin) with strong mechanistic ties to HPV-positive HNC biology.

**Table A4.**
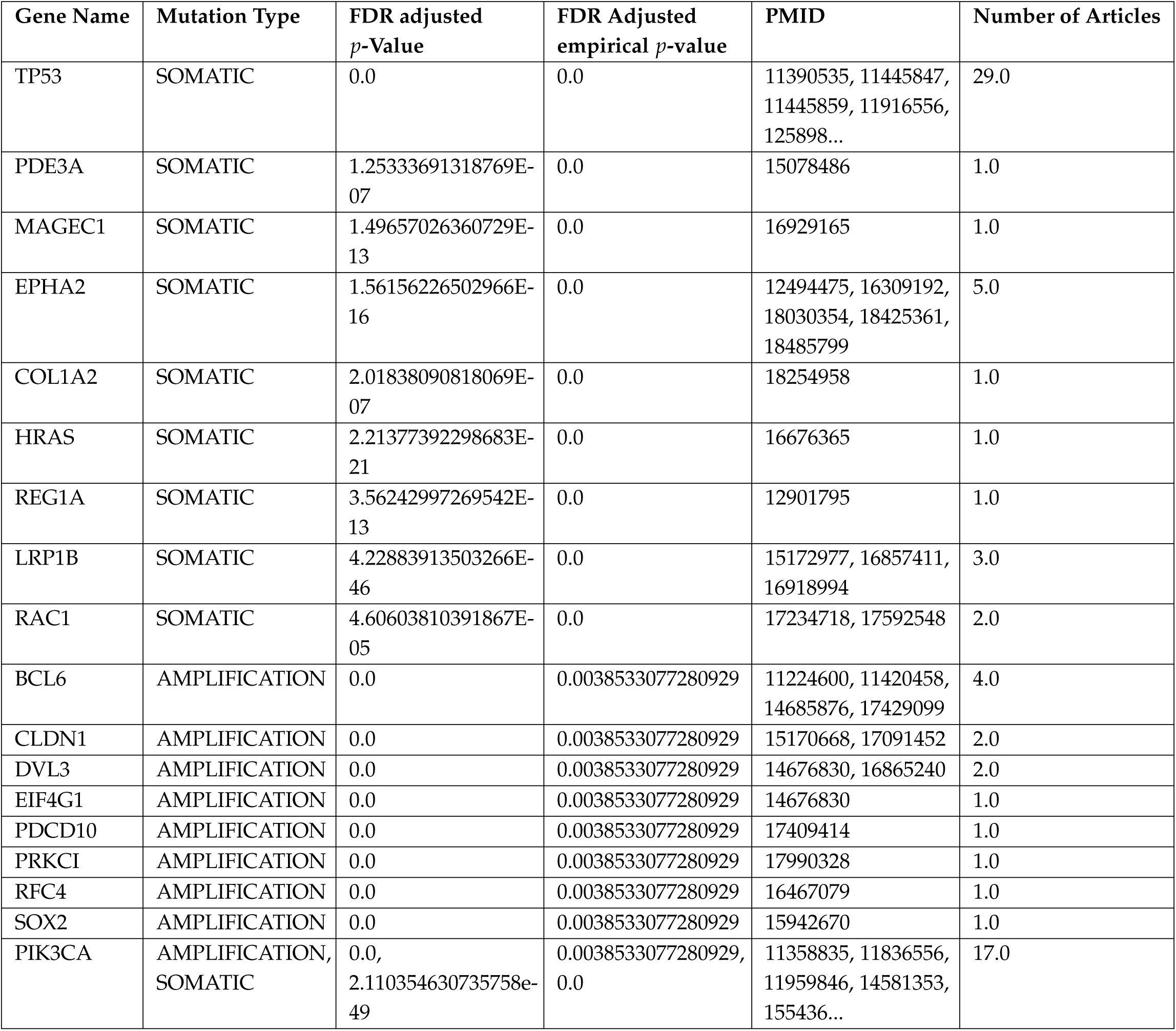
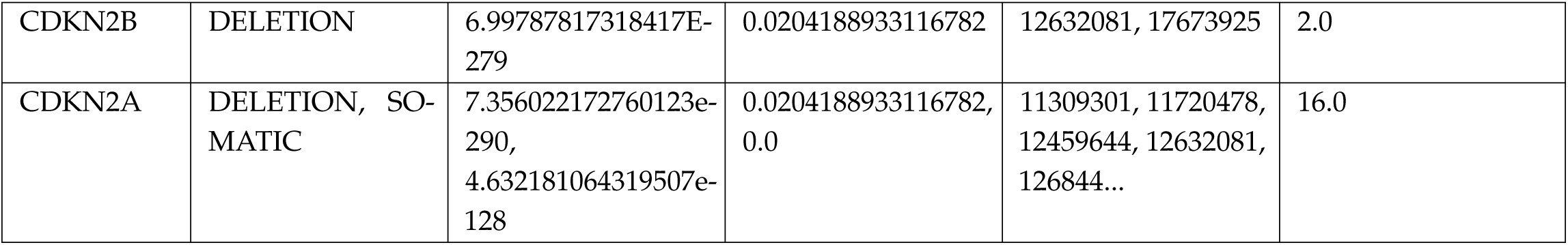
Top High-Risk Genes Identified in HPV-Negative HNC Cohort. This figure highlights the most significant genes identified through integrative analysis of copy number variation (CNV) and somatic mutation profiles in HPV-negative tumors. Genes such as SOX2, PRKCI, EPHA2, PIK3CA and TP53 emerged as key oncogenic drivers, alongside pathway-level regulators like EIF4G1 and CLDN1. All listed genes demonstrated strong statistical significance (FDR adjusted *p*-value < 1.25 E-7; FDR adjusted empirical *p*-value 0.003858), indicating recurrent amplifications or somatic mutations across the cohort. These targets underpin critical biological processes, including chromatin remodeling, signal transduction, and immune modulation, and were prioritized for downstream drug repurposing analysis.

**Table A5.** Direct drug–gene associations identified for the HPV-negative HNC cohort are summarized in this table. Columns include drug name, literature-validated gene targets, mutation type (MUT TYPE), supporting PMID references, number of supporting articles, and empirical FDR-adjusted *p*-values. These medications directly target high-confidence genomic alterations such as amplifications in PRKCI, and PIK3CA, as well as somatic mutations in TP53 and EPHA2, which define the molecular landscape of HPV-negative tumors. All gene targets were validated through a large-scale literature mining pipeline to ensure biological relevance, and only candidates with confirmed associations to HNC were retained. Literature validation strengthens mechanistic confidence, while empirical FDR adjustment ensures statistical robustness. These direct associations informed prioritization of top candidates, including multi-kinase inhibitors (Fostamatinib, Regorafenib) and pathway modulators (Acetylsalicylic acid), which emerged as promising repurposing opportunities for precision therapy in HPV-negative HNC.

| DRUG | Drug FDR adjusted Empirical *p*-value | Mutation Type | Literature-Validated Gene Targets | Article Count | PMIDs | Drug-Gene Interaction |
| --- | --- | --- | --- | --- | --- | --- |
| Bisindolylmaleimide I | 0.002927 | AMPLIFICATION | PRKCI | 1 | 17990328 | PRKCI: UNKNOWN |
| Fostamatinib | 0.002927 | AMPLIFICATION, SOMATIC | EPHA2, PRKCI | 6 | 17990328, 18030354, 16309192, 18425361, 124944... | EPHA2: inhibitor; PRKCI: inhibitor |
| Acetylsalicylic acid | 0.004258 | SOMATIC | TP53 | 29 | 17638058, 11916556, 16969480, 16682429, 173127... | TP53: inducer |
| Dasatinib | 0.004258 | SOMATIC | EPHA2 | 5 | 18030354, 16309192, 18425361, 12494475, 18485799 | EPHA2: antagonist |
| Regorafenib | 0.004258 | SOMATIC | EPHA2 | 5 | 18030354, 16309192, 18425361, 12494475, 18485799 | EPHA2: inhibitor |
| Wortmannin | 0.046275 | AMPLIFICATION, SOMATIC | PIK3CA | 17 | 16380997, 11358835, 14581353, 15700036, 168070... | PIK3CA: UNKNOWN |
| XL765 | 0.046301 | AMPLIFICATION,<br>SOMATIC | PIK3CA | 17 | 16380997,<br>11358835,<br>14581353,<br>15700036,<br>168070... | PIK3CA: UNKNOWN |

**Table A6.** Top 50 indirect drug–gene associations for the HPV-negative HNC cohort, ranked by empirical FDR-adjusted *p*-values. Columns include drug name, literature-validated gene targets, associated connected risk gene, mutation type (MUT TYPE), supporting PMID references, number of supporting articles, and empirical FDR-adjusted *p*-values. These medications target immediate network neighbors of high-confidence risk genes identified through CNV and somatic mutation analyses, leveraging STRING PPI data to capture pathway-level relevance. All gene targets and their network neighbors were validated through a large-scale literature mining pipeline to ensure biological significance, and only candidates with confirmed associations to HNC were retained. This network-based expansion broadens the therapeutic landscape beyond direct genomic alterations, uncovering multi-kinase inhibitors (e.g., Brigatinib and Regorafenib) and novel agents (Fostamatinib, Artenimol, Quercetin) with strong mechanistic ties to HPV-negative HNC biology. Literature validation and statistical prioritization reinforce the translational potential of these candidates for precision drug repurposing.

| DRUG | Drug Empirical FDR | Literature-Validated Gene Targets | Literature-Validated Risk Genes | Risk Gene Mutation Type | Gene Target Article Count | Gene Target PMIDs | Risk Gene Article Count | Risk Gene PMIDs | Drug-Gene Interaction |
| --- | --- | --- | --- | --- | --- | --- | --- | --- | --- |
| brigatinib | 0.004258 | ALK, EGFR, ERBB2, ERBB4, IGF1R, MET | CDKN2A, PIK3CA | AMPLIFICATION, DELETION, SOMATIC | 1004 | 17826531, 14603452, 15161032, 17250463, 170984... | 50 | 15495191, 18413807, 16807070, 11309301, 168496... | ALK: inhibitor; EGFR: inhibitor; ERBB2: inhibi... |
| erdafitinib | 0.004258 | FGFR1, FGFR2, FGFR3, FGFR4, KIT, PDGFRA, RET | PIK3CA | AMPLIFICATION, SOMATIC | 134 | 11329134, 16807070, 16342249, 11953175, 172531... | 34 | 16380997, 11358835, 14581353, 15700036, 168070... | FGFR1: inhibitor; FGFR2: inhibitor; FGFR3: inh... |
| lenvatinib | 0.004258 | FGFR1, FGFR2, FGFR3, FGFR4, KIT, PDGFRA, RET | PIK3CA | AMPLIFICATION, SOMATIC | 134 | 11329134, 16807070, 16342249, 11953175, 172531... | 34 | 16380997, 11358835, 14581353, 15700036, 168070... | FGFR1: inhibitor; FGFR2: inhibitor; FGFR3: inh... |
| lucitanib | 0.004258 | FGFR1, FGFR2, FGFR3, PDGFRA | PIK3CA | AMPLIFICATION, SOMATIC | 24 | 17545628, 16807070, 11562460, 11605053, 175135... | 34 | 16380997, 11358835, 14581353, 15700036, 168070... | FGFR1: inhibitor; FGFR2: inhibitor; FGFR3: inh... |
| nintedanib | 0.004258 | FGFR1, FGFR2, FGFR3, LYN, PDGFRA, SRC | PIK3CA | AMPLIFICATION, SOMATIC | 28 | 17545628, 16807070, 11562460, 17252232, 116050... | 34 | 16380997, 11358835, 14581353, 15700036, 168070... | FGFR1: inhibitor; FGFR2: inhibitor; FGFR3: inh... |
| pd-166326 | 0.004258 | EGFR, FGFR1, KIT, PDGFRA, SRC | CDKN2A, PIK3CA, TP53 | AMPLIFICATION, DELETION, SOMATIC | 921 | 17671674, 17964995, 12440226, 17826531, 146034... | 79 | 15495191, 18413807, 11916556, 16807070, 176380... | EGFR: inhibitor; FGFR1: inhibitor; KIT: inhibi... |
| sorafenib | 0.004258 | BRAF, EGFR, FGFR1, KIT, RET | PIK3CA | AMPLIFICATION, SOMATIC | 790 | 17826531, 14603452, 15161032, 17250463, 170984... | 34 | 16380997, 11358835, 14581353, 15700036, 168070... | BRAF: inhibitor; EGFR: inhibitor; FGFR1: inhib... |
| sunitinib | 0.004258 | KIT, MET, PDGFRA | PIK3CA | AMPLIFICATION, SOMATIC | 40 | 16342249, 17253141, 11911252, 11705871, 170751... | 34 | 16380997, 11358835, 14581353, 15700036, 168070... | KIT: inhibitor; MET: inhibitor; PDGFRA: inhibitor |
| ponatinib | 0.004258 | FGFR1, FGFR2, FGFR3, FGFR4, KIT, LYN, PDGFRA, ... | CDKN2A, PIK3CA | AMPLIFICATION, DELETION, SOMATIC | 144 | 11329134, 16807070, 16342249, 17961551, 119531... | 50 | 15495191, 18413807, 16807070, 11309301, 168496... | FGFR1: inhibitor; FGFR2: inhibitor; FGFR3: inh... |
| pralsetinib | 0.004258 | FGFR1, FGFR2, JAK2, NTRK1, NTRK3, RET | PIK3CA | AMPLIFICATION, SOMATIC | 118 | 11329134, 16807070, 11279608, 11953175, 159883... | 34 | 16380997, 11358835, 14581353, 15700036, 168070... | FGFR1: inhibitor; FGFR2: inhibitor; JAK2: inhi... |
| pp-121 | 0.004258 | EGFR, HCK, MTOR, PDGFRA, SRC | CDKN2A, PIK3CA, TP53 | AMPLIFICATION, DELETION, SOMATIC | 915 | 17826531, 14603452, 15161032, 17250463, 170984... | 79 | 15495191, 18413807, 11916556, 16807070, 176380... | EGFR: inhibitor; HCK: inhibitor; MTOR: inhibit... |
| regorafenib | 0.004258 | BRAF, EPHA2, FGFR1, FGFR2, KIT, NTRK1, PDGFRA,... | CLDN1, PIK3CA, TP53 | AMPLIFICATION, SOMATIC | 217 | 11329134, 17638058, 17097223, 14508525, 129076... | 65 | 17638058, 11916556, 16807070, 16969480, 178483... | BRAF: inhibitor; EPHA2: inhibitor; FGFR1: inhi... |
| acetylsalicylic acid | 0.004258 | CCND1, MYC, NFKBIA, PCNA, PTGS2, TP53 | HRAS, PIK3CA, TP53 | AMPLIFICATION, SOMATIC | 114 | 17638058, 11854069, 12854173, 11173150, 110706... | 47 | 17638058, 11916556, 16807070, 16969480, 178483... | CCND1: downregulator; MYC: downregulator; NFKB... |
| bisindolylmaleimide i | 0.004258 | AKT1, CDK1, CHEK1, MAPK8, PRKCI | CDKN2A, PIK3CA, RFC4, SOX2, TP53 | AMPLIFICATION, DELETION, SOMATIC | 35 | 16525614, 15896313, 14581353, 15645429, 168886... | 83 | 15495191, 18413807, 11916556, 16807070, 176380... | AKT1: inhibitor; CDK1: inhibitor; CHEK1: inhib... |
| dasatinib | 0.004258 | EPHB4, FYN, HSPA8, KIT, LYN, SRC, STAT5B, YES1 | EPHA2, HRAS, PIK3CA, TP53 | AMPLIFICATION, SOMATIC | 18 | 17611172, 16342249, 11973644, 12771142, 117562... | 52 | 17638058, 11916556, 16807070, 16969480, 178483... | EPHB4: inhibitor; FYN: inhibitor; HSPA8: binde... |
| fostamatinib | 0.004258 | ALK, AURKA, BRAF, CDK1, CDK4, CHEK1, DAPK1, EG... | CLDN1, EIF4G1, EPHA2, HRAS, PIK3CA, RAC1, RFC4... | AMPLIFICATION, SOMATIC | 1008 | 11854069, 17826531, 14603452, 15161032, 150156... | 80 | 17638058, 11916556, 16807070, 16969480, 178483... | ALK: inhibitor; AURKA: inhibitor; BRAF: inhibi... |
| pazopanib | 0.004930 | FGFR3, KIT, PDGFRA | PIK3CA | AMPLIFICATION, SOMATIC | 24 | 16342249, 11605053, 17317803, 17253141, 175135... | 39 | 16807070, 17848307, 16676365, 17990317, 163809... | FGFR3: inhibitor; KIT: inhibitor; PDGFRA: inhi... |
| quercetin | 0.004930 | ESR1, HCK, STAT3 | PIK3CA | AMPLIFICATION, SOMATIC | 44 | 17145757, 17525119, 17826306, 16490172, 166424... | 17 | 16380997, 11358835, 14581353, 15700036, 168070... | ESR1: UNKNOWN; HCK: UNKNOWN; STAT3: inhibitor |
| fn-1501 | 0.006245 | CDK2, CDK4, CDK6 | CDKN2A, CDKN2B | DELETION, SOMATIC | 33 | 12017338, 11267947, 11453659, 11854069, 173401... | 18 | 15495191, 16713672, 16278815, 12632081, 184138... | CDK2: inhibitor; CDK4: inhibitor; CDK6: inhibitor |
| olomoucine | 0.006245 | CDK1, CDK2, CDK5 | CDKN2A, CDKN2B | DELETION, SOMATIC | 16 | 16525614, 17145757, 11267947, 11453659, 156454... | 18 | 15495191, 16713672, 16278815, 12632081, 184138... | CDK1: binder; CDK2: inhibitor; CDK5: inhibitor |
| purvalanol | 0.006245 | CDK1, CDK2, CDK4, CDK5 | CDKN2A, CDKN2B | DELETION, SOMATIC | 30 | 16525614, 17145757, 11267947, 11453659, 120173... | 18 | 15495191, 16713672, 16278815, 12632081, 184138... | CDK1: binder; CDK2: inhibitor; CDK4: UNKNOWN; ... |
| tetrabromo-2-benzotriazole | 0.006855 | AKT1, CHEK1, MAPK8 | CDKN2A, PIK3CA, RFC4, TP53 | AMPLIFICATION, DELETION, SOMATIC | 24 | 15896313, 14581353, 17974918, 12532415, 152266... | 82 | 15495191, 18413807, 11916556, 16807070, 176380... | AKT1: inhibitor; CHEK1: inhibitor; MAPK8: inhi... |
| seliciclib | 0.006939 | CDK1, CDK2 | TP53 | SOMATIC | 15 | 16525614, 12017338, 11267947, 11453659, 156454... | 46 | 17638058, 11916556, 16807070, 16969480, 178483... | CDK1: inhibitor; CDK2: inhibitor |
| ci-1040 | 0.006939 | AKT1, CHEK1, MAPK8 | CDKN2A, PIK3CA, RFC4, TP53 | AMPLIFICATION, DELETION, SOMATIC | 24 | 15896313, 14581353, 17974918, 12532415, 152266... | 82 | 15495191, 18413807, 11916556, 16807070, 176380... | AKT1: inhibitor; CHEK1: inhibitor; MAPK8: inhi... |
| resveratrol | 0.006939 | AKT1, APP, ESR1, ITGB3, PTGS2 | COL1A2, PIK3CA, TP53 | AMPLIFICATION, SOMATIC | 10 | 15896313, 14581353, 11309301, 16458426, 179749... | 47 | 17638058, 11916556, 16807070, 16969480, 178483... | AKT1: inhibitor; APP: UNKNOWN; ESR1: UNKNOWN; ... |
| arsenic trioxide | 0.006939 | AKT1, CCND1, CDKN1A, HDAC1, PML | CDKN2A, CDKN2B, PIK3CA, TP53 | AMPLIFICATION, DELETION, SOMATIC | 67 | 15896313, 18398822, 16807070, 16446704, 125324... | 64 | 15495191, 18413807, 17673925, 11916556, 176380... | AKT1: inducer; CCND1: antagonist; CDKN1A: UNKN... |
| midostaurin | 0.008516 | KIT, PDGFRA | PIK3CA | AMPLIFICATION, SOMATIC | 18 | 16342249, 17317803, 17253141, 17513510, 144996... | 34 | 16380997, 11358835, 14581353, 15700036, 168070... | KIT: antagonist; PDGFRA: antagonist |
| semaxanib | 0.008516 | FGFR1, KIT, PDGFRA, RET | PIK3CA | AMPLIFICATION, SOMATIC | 61 | 11329134, 16807070, 16342249, 11953175, 172531... | 17 | 16380997, 11358835, 14581353, 15700036, 168070... | FGFR1: inhibitor; KIT: inhibitor; PDGFRA: inhi... |
| hymenialdisine | 0.010409 | CDK1, CDK2, CDK5 | CDKN2A, CDKN2B | DELETION, SOMATIC | 16 | 16525614, 17145757, 11267947, 11453659, 156454... | 18 | 15495191, 16713672, 16278815, 12632081, 184138... | CDK1: UNKNOWN; CDK2: inhibitor; CDK5: inhibitor |
| su9516 | 0.010409 | CDK1, CDK2, CDK5 | CDKN2A, CDKN2B | DELETION, SOMATIC | 16 | 16525614, 17145757, 11267947, 11453659, 156454... | 18 | 15495191, 16713672, 16278815, 12632081, 184138... | CDK1: binder; CDK2: inhibitor; CDK5: inhibitor |
| martinostat | 0.013952 | HDAC1 | BCL6 | AMPLIFICATION | 1 | 12239610 | 4 | 17429099, 11420458, 11224600, 14685876 | HDAC1: inhibitor |
| tivozanib | 0.016152 | FGFR1, KIT, MET, PDGFRA | PIK3CA | AMPLIFICATION, SOMATIC | 50 | 16807070, 16342249, 17253141, 11911252, 117058... | 34 | 16380997, 11358835, 14581353, 15700036, 168070... | FGFR1: inhibitor; KIT: inhibitor; MET: inhibit... |
| ripretinib | 0.016532 | BRAF, KIT, PDGFRA | PIK3CA | AMPLIFICATION, SOMATIC | 49 | 16728576, 17638058, 17097223, 16342249, 182515... | 17 | 16380997, 11358835, 14581353, 15700036, 168070... | BRAF: inhibitor; KIT: inhibitor; PDGFRA: inhib... |
| famitinib | 0.018131 | KIT, PDGFRA | PIK3CA | AMPLIFICATION, SOMATIC | 18 | 16342249, 17317803, 17253141, 17513510, 144996... | 34 | 16380997, 11358835, 14581353, 15700036, 168070... | KIT: inhibitor; PDGFRA: inhibitor |
| artenimol | 0.018131 | ANXA2, GAPDH, HSPA8, LGALS1, PRDX1, RPS13, RPS... | EIF4G1, HRAS, SOX2, TP53 | AMPLIFICATION, SOMATIC | 24 | 17540725, 15381369, 15311506, 17565818, 163167... | 85 | 15495191, 18413807, 11916556, 16807070, 176380... | ANXA2: ligand; GAPDH: ligand; GAPDH: ligand; H... |
| ag-24322 | 0.018736 | CDK1, CDK2, CDK4 | CDKN2A, CDKN2B | DELETION, SOMATIC | 29 | 16525614, 12017338, 11267947, 11453659, 118540... | 18 | 15495191, 16713672, 16278815, 12632081, 184138... | CDK1: inhibitor; CDK2: inhibitor; CDK4: inhibitor |
| mkc-1 | 0.018736 | CDK1, CDK2, CDK4 | CDKN2A, CDKN2B | DELETION, SOMATIC | 29 | 16525614, 12017338, 11267947, 11453659, 118540... | 18 | 15495191, 16713672, 16278815, 12632081, 184138... | CDK1: inhibitor; CDK2: inhibitor; CDK4: inhibitor |
| pd-168393 | 0.018736 | EGFR, ERBB2, SRC | CDKN2A | DELETION, SOMATIC | 321 | 17826531, 14603452, 15161032, 17250463, 170984... | 16 | 15495191, 16713672, 16278815, 12632081, 184138... | EGFR: inhibitor; ERBB2: inhibitor; SRC: UNKNOWN |
| pha-793887 | 0.018736 | CDK1, CDK2, CDK4 | CDKN2A, CDKN2B | DELETION, SOMATIC | 29 | 16525614, 12017338, 11267947, 11453659, 118540... | 18 | 15495191, 16713672, 16278815, 12632081, 184138... | CDK1: inhibitor; CDK2: inhibitor; CDK4: inhibitor |
| enzastaurin | 0.020074 | AKT1, AURKA, CHEK1 | CDKN2A, PIK3CA, TP53 | AMPLIFICATION, DELETION, SOMATIC | 22 | 15896313, 14581353, 18311783, 17974918, 125324... | 79 | 15495191, 18413807, 11916556, 16807070, 176380... | AKT1: inhibitor; AURKA: UNKNOWN; CHEK1: UNKNOWN |
| prasterone | 0.024089 | AR, ESR1 | PIK3CA, TP53 | AMPLIFICATION, SOMATIC | 11 | 16376136, 11556855, 18528886, 18429833, 128174... | 46 | 17638058, 11916556, 16807070, 16969480, 178483... | AR: agonist; ESR1: binder |
| trilaciclib | 0.024089 | CDK2, CDK4,<br>CDK5, CDK6 | CDKN2A,<br>CDKN2B, PIK3CA,<br>TP53 | AMPLIFICATION,<br>DELETION, SO-MATIC | 68 | 17145757,<br>12017338,<br>11267947,<br>11453659,<br>118540... | 64 | 15495191,<br>18413807,<br>17673925,<br>11916556,<br>176380... | CDK2: inhibitor;<br>CDK4: inhibitor;<br>CDK5: inhibi... |
| selpercatinib | 0.028624 | FGFR1, FGFR2,<br>FGFR3, RET | PIK3CA | AMPLIFICATION,<br>SOMATIC | 114 | 11329134,<br>16807070,<br>11953175,<br>15988378,<br>155838... | 34 | 16380997,<br>11358835,<br>14581353,<br>15700036,<br>168070... | FGFR1: inhibitor;<br>FGFR2: inhibitor;<br>FGFR3: inh... |
| amuvatinib | 0.028824 | KIT, MET,<br>PDGFRA, RAD51,<br>RET | PIK3CA, TP53 | AMPLIFICATION,<br>SOMATIC | 136 | 11329134,<br>16342249,<br>11953175,<br>17253141,<br>159883... | 63 | 17638058,<br>11916556,<br>16807070,<br>16969480,<br>178483... | KIT: modulator;<br>MET: UNKNOWN;<br>PDGFRA: modu-<br>lato... |
| linifanib | 0.028824 | KIT | PIK3CA | AMPLIFICATION,<br>SOMATIC | 14 | 16342249,<br>17317803,<br>17253141,<br>17513510,<br>179352... | 34 | 16380997,<br>11358835,<br>14581353,<br>15700036,<br>168070... | KIT: UNKNOWN |
| xl999 | 0.028824 | FGFR1, FGFR3,<br>RET | PIK3CA | AMPLIFICATION,<br>SOMATIC | 55 | 11329134,<br>16807070,<br>11953175,<br>15988378,<br>155838... | 17 | 16380997,<br>11358835,<br>14581353,<br>15700036,<br>168070... | FGFR1: UN-KNOWN; FGFR3:<br>UNKNOWN; RET:<br>UNKNOWN |
| canertinib | 0.028824 | AKT1, EGFR,<br>ERBB2 | CDKN2A | DELETION, SO-MATIC | 326 | 17826531,<br>14603452,<br>15161032,<br>17250463,<br>170984... | 16 | 15495191,<br>16713672,<br>16278815,<br>12632081,<br>184138... | AKT1: inhibitor;<br>EGFR: inhibitor;<br>ERBB2: inhib... |
| alsterpaullone | 0.030111 | CDK1, CDK2,<br>CDK5 | CDKN2A,<br>CDKN2B | DELETION, SO-MATIC | 16 | 16525614,<br>17145757,<br>11267947,<br>11453659,<br>156454... | 18 | 15495191,<br>16713672,<br>16278815,<br>12632081,<br>184138... | CDK1: inhibitor;<br>CDK2: inhibitor;<br>CDK5: inhibitor |
| zanubrutinib | 0.030714 | EGFR, ERBB2, ERBB4, JAK2, JAK3 | PIK3CA | AMPLIFICATION, SOMATIC | 327 | 17826531, 14603452, 15161032, 17250463, 170984... | 17 | 16380997, 11358835, 14581353, 15700036, 168070... | EGFR: inhibitor; ERBB2: inhibitor; ERBB4: inhi... |
| foreskin keratinocyte (neonatal) | 0.037918 | EGFR, FGFR2, IL1B, IL6, PDGFRA, TNF | EPHA2, PIK3CA, TP53 | AMPLIFICATION, SOMATIC | 308 | 17826531, 14603452, 15161032, 17250463, 170984... | 51 | 17638058, 11916556, 16807070, 16969480, 178483... | EGFR: agonist; FGFR2: agonist; IL1B: agonist; ... |

## Disclaimer/Publisher’s Note

The statements, opinions and data contained in all publications are solely those of the individual author(s) and contributor(s) and not of MDPI and/or the editor(s). MDPI and/or the editor(s) disclaim responsibility for any injury to people or property resulting from any ideas, methods, instructions or products referred to in the content.

